# Landscape context drives within-species variation in dispersal kernels and limits transferability

**DOI:** 10.1101/2025.11.09.687206

**Authors:** Jette Wolff, Guillermo Fandos, Cécile H. Albert, Greta Bocedi, Justin M. J. Travis, Damaris Zurell

## Abstract

Dispersal is a key ecological trait that ensures connectivity, gene flow, and range dynamics, yet empirical information about the dispersal process remains scarce. While interspecific differences have been synthesized, intraspecific variation driven by individual movement and landscape context remains poorly understood. Here, we use the individual-based modelling platform *RangeShifter* to simulate movement trajectories of four animal species, from insects to mammals, across neutral landscapes varying in habitat amount and fragmentation. From these simulations we derived dispersal kernels using five probability density functions and analysed their response to landscape context. Across species, log-normal (fat-tailed) kernels consistently best described dispersal, but both median and long-distance dispersal varied strongly with habitat configuration. Habitat loss consistently reduced abundances, whereas fragmentation per se more strongly shaped dispersal kernels, providing mechanistic insight into a central debate in landscape ecology. Our results highlight that dispersal kernels do not transfer reliably across environments and should be treated as emergent, context-dependent properties. Progress in predicting species’ responses to global change will require empirical studies explicitly integrated with mechanistic modelling to uncover generalizable links between dispersal, traits, and landscape context.

## Introduction

Dispersal is a crucial ecological trait that ensures connectivity between populations, drives gene flow, and influences range dynamics (Bonte and Dahirel 2017). It also has significant implications for biodiversity patterns, ecosystem dynamics, and the effectiveness of conservation strategies. A deep understanding of the mechanisms underpinning dispersal is essential for predicting changes in biodiversity, especially in the face of environmental and land use changes. This knowledge is crucial for anticipating how species will respond to altered landscapes, guiding targeted conservation efforts and informed land-use planning. Traditionally, dispersal distances are modelled using dispersal kernels, which depict the density distribution of individuals traveling a given distance (Hovestadt et al. 2012, Nathan et al. 2012). However, these models rely on scarce empirical data, resulting in high levels of uncertainty and an incomplete understanding of dispersal dynamics. This constrains the accuracy of predictions, highlighting the need for improved datasets and more robust approaches to better capture the complexity of dispersal processes (e.g. Chadœuf et al. 2017).

More precisely, dispersal encompasses individual movements from natal to breeding patches (natal dispersal) or between consecutive breeding patches (breeding dispersal). Dispersal consists of three distinct phases: emigration, transfer, and settlement (Clobert et al. 2012). Emigration is often influenced by density-dependent factors (Matthysen 2005, Jreidini and Green 2024), serving as a mechanism to avoid inbreeding when density is low, or to mitigate kin and intra-specific competition for resources when density is high. The transfer phase involves the actual movement through the landscape, where various environmental elements influence movement trajectories. Finally, settlement may depend on habitat suitability, population density and the presence of potential mates. Each of these phases is associated with potential costs to the dispersing individual (Bonte et al. 2012).

Quantifying these phases across spatial scales is challenging, as it requires detailed tracking of movement patterns and of rare events. New tracking technologies have greatly improved empirical analyses allowing in-depth study of these dispersal phases. Yet, detailed movement data are typically confined to a limited number of individuals, which poses challenges in accurately capturing dispersal per se (beyond daily movements and potential dispersal attempts) and in particular rarer long-distance dispersal events (LDD). Though rarer, LDD events are pivotal for species adapting to shifting environmental conditions and colonizing new habitats (Nathan et al. 2003, 2012, Nathan 2006, Gillespie et al. 2012, Jordano 2017) LDD events are better captured in empirical dispersal kernels that provide information about the complete set of dispersal distances observable across a population (Bullock et al. 2017). Typically, seed-trap or mark-recapture (including bird ringing) data are used to inversely fit mathematical kernels to dispersal distances observed across many individuals, which increases the probability to observe the rare LDD events (Nathan et al. 2012, Rogers et al. 2019).

The study of dispersal has a long history, with early efforts focusing on collating dispersal distances and making allometric inferences across different species (Sutherland et al. 2000, Bowman et al. 2002, Stevens et al. 2013, Whitmee and Orme 2013). More recently, empirical dispersal kernels have been synthesized for plants (Bullock et al. 2017) and birds (Fandos et al. 2023). Together, these studies provide valuable information for making accurate predictions about population connectivity and range dynamics in conservation and climate impact assessments, at least for a few very well tracked systems and species. However, these distances and kernels estimate often remain largely phenomenological, focused on inter-specific differences, and may be limited in their transferability to different populations or geographic regions (Cote et al. 2017). Understanding the mechanistic underpinnings of these empirical patterns requires examining how individual movement processes scale up to population-level dispersal kernels.

Despite these ancient and work-intensive efforts to synthesize knowledge on dispersal distances and kernels among species and groups, we still know little about what will shape variation in dispersal kernels and distances within species. According to the movement ecology paradigm (Nathan 2008), individual movement paths are determined by intrinsic factors, such as motion and navigation capacity, and external factors related to the biotic and abiotic environment and stochasticity. Phenotypic differences between dispersing individuals and differences in the landscape context may modify the dispersal outcome and the probability of LDD (Clobert et al. 2009, Bonte and Dahirel 2017). Specifically, landscape configuration, including the amount and spatial arrangement of suitable habitat, as well as the degree of habitat fragmentation and isolation related to matrix unsuitability, influence the transfer process by mediating movement costs, mortality risks and barrier effects (Argote-Deluque et al. 2025).

Understanding how landscape configuration shapes dispersal connects directly to the long-standing debate on the relative importance of habitat loss versus fragmentation per se (Fahrig 2003, Haddad et al. 2015). Dispersal provides a key mechanistic link in this debate, as movement processes may respond differently to changes in habitat amount versus spatial arrangement. Studies have demonstrated that landscape dynamics, the cost of dispersal, and the level of environmental stochasticity affect the correlation between dispersal and other morphological, physiological, and behavioural traits, referred to as dispersal syndromes (Ronce et al. 2000, Bonte and De La Peña 2009, Clobert et al. 2009, Bonte and Dahirel 2017, Cote et al. 2022). Improving our understanding of how dispersal kernels emerge from individual movement decisions depending on environmental contexts is vital for predicting dispersal and, consequently, species responses to changing environmental conditions.

Studying the context dependency of dispersal is challenged by data availability as it requires movement or dispersal information across many individuals in different landscape contexts. Such data are difficult to obtain or limited to few taxonomic groups or species. For example, bird ringing data allow estimating dispersal distances across many bird species, but different biases in resighting or recapture of different bird species result in uneven sample sizes across species (Chu and Claramunt 2023, Fandos et al. 2023). Mark-recapture studies are also common for studying butterfly dispersal but are often limited to a few well-studied species (Stevens et al. 2010). Finally, tracking data provide perhaps the most detailed information on the dispersal process but are typically restricted to larger bodied species and vertebrates, and to comparably few individuals, leading to a knowledge shortfall in species movements (Scarpignato et al. 2023). Theoretical models provide a useful alternative for studying dispersal in different landscape contexts e.g.(Travis and Dytham 1999, Fahrig 2007). Individual-based modelling (IBM) approaches are particularly well suited to understand how dispersal emerges from individual movement decisions and environmental context (Palmer et al. 2011, Simpkins et al. 2018, Bauduin et al. 2020). Previous theoretical studies have provided important insights on the role of habitat fragmentation on range expansion (Bocedi et al. 2014), the costs of dispersal (Schtickzelle et al. 2006) and the evolution of dispersal rates (Travis and Dytham 1999). However, the potential of theoretical models for understanding emerging dispersal patterns across different landscape contexts and taxonomic groups remains untapped.

Here, we aim to understand how landscape context influences emerging dispersal patterns. Using the individual-based modelling platform RangeShifter (Bocedi et al. 2021, Malchow et al. 2021) to simulate individual-level movement processes across landscapes scenarios, we investigate the impact of different degrees of landscape fragmentation and habitat amount on the shape of population-level dispersal kernels as well as key dispersal measures. Our simulations encompass dispersal and population dynamics of four different taxonomic groups ranging from small insects to large mammals based on case studies previously parameterized in RangeShifter (Aben et al. 2016, Ovenden et al. 2019, Melero et al. 2020, Williams et al. 2021). Simulations are set within neutral landscapes varying in the degree of fragmentation (from single large to several small habitat patches) and in the total amount of available habitat. From each simulation, we extracted the start and end point of the dispersal process from individual movement paths (thus representing tracking data) and estimated dispersal kernels using different probability density functions that vary in their shape and their ability to capture long-distance dispersal (Fandos et al. 2023, Figure 1). We then analysed how median and long-distance dispersal, defined as 50% and 97.5% quantiles of the probability density functions, varied across different levels of habitat fragmentation and habitat amount.

**Figure 1:**
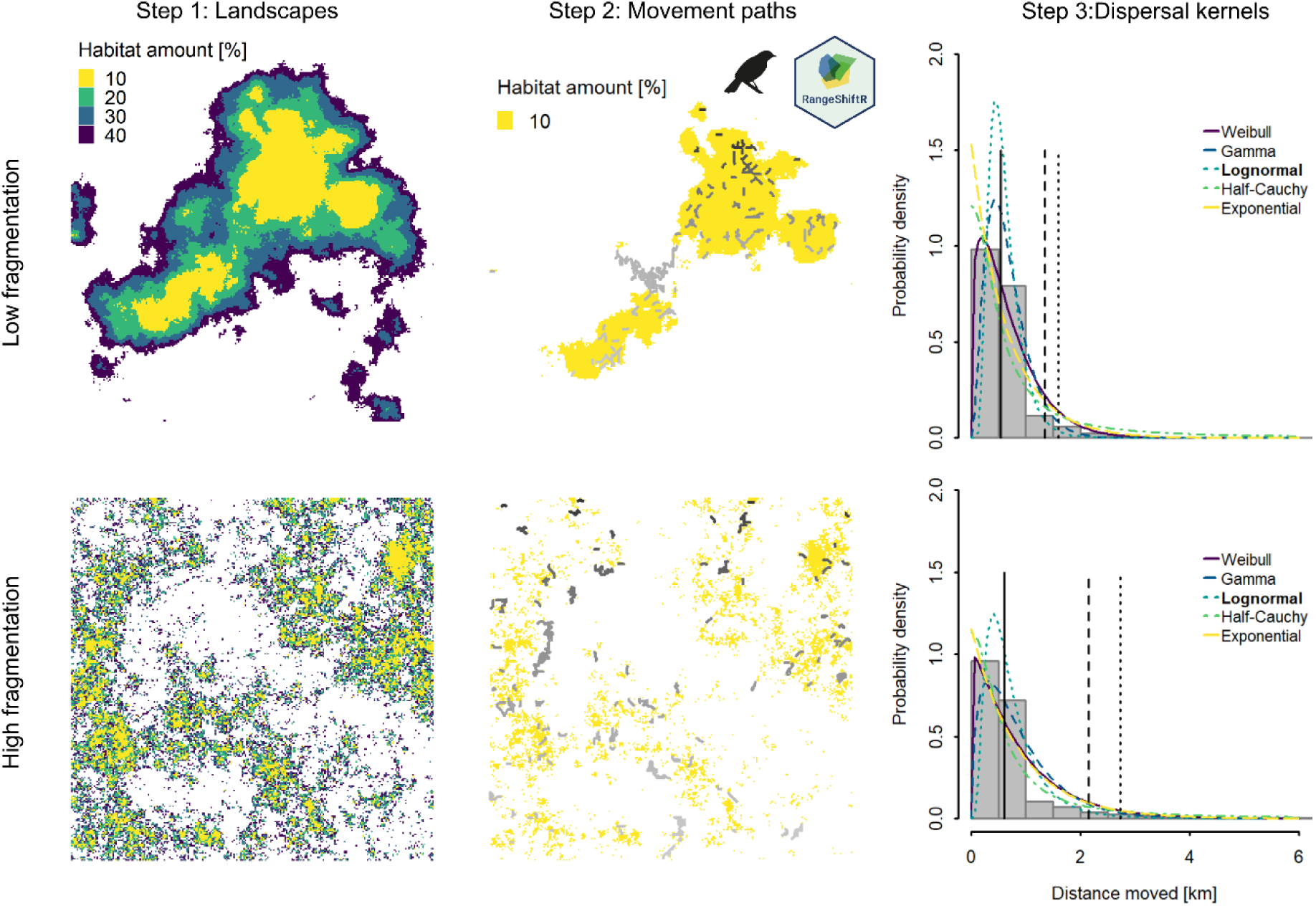
Workflow of the simulations and statistical analyses. Step 1 (left panels): We generated landscapes with different landscape configurations (varying amount of habitat and degree of fragmentation). Landscapes were generated using the midpoint displacement algorithms with low and high fragmentation corresponding to low and high roughness parameters, respectively. Step 2 (central panels): Using four previously published RangeShifter case studies, we simulated movement paths of individuals. Movement paths of the bird case in one landscape repetition with 10% habitat amount are represented in shades of grey (for 100 randomly sampled individuals). Step 3 (right panel, same x-axis for both graphs): We calculated the corresponding distances between the start and endpoint of dispersal for all successful dispersers – mimicking mark-recapture data - and estimated the parameters for five different types of dispersal kernels. The vertical lines represent the median (solid line), 97.5^th^ (dashed line) and 99^th^ (dotted line) quantiles based on the kernel with the best fit (marked in bold).

The effect of habitat amount and fragmentation is likely to be complex and depending on species life-histories and dispersal mode) (Cote et al. 2017), making it challenging to formulate universal predictions. Nevertheless, we make two key predictions based on movement ecology theory. First, we predict that higher habitat amount will reduce dispersal distances due to the “shadow effect” whereby abundant nearby patches provide immediate settlement opportunities (Hein et al. 2004, Heinz et al. 2005, Bocedi et al. 2014). Second, we predict that fragmentation effects will depend on species-specific matrix resistance. Species with low matrix resistance should disperse farther in fragmented landscapes as they successfully traverse the matrix to reach distant suitable patches. Conversely, species with high matrix resistance should show reduced dispersal distances in fragmented landscapes, for example through increased mortality and movement costs that limit their ability and motivation to cross the matrix (Fletcher Jr. et al. 2019).

## Methods

### Landscapes

We employed a neutral landscape generator (Sciaini et al. 2018) to create landscapes with varying degrees of fragmentation using the midpoint displacement algorithm. This algorithm incorporates a roughness parameter *r,* which controls the degree of spatial autocorrelation within the landscape ranging from 0 to 1. A roughness value of 1 corresponds to the lowest spatial autocorrelation, indicative of high fragmentation with many small habitat patches. We selected roughness values from 0.55 to 0.95 in increments of 0.1, resulting in five levels of roughness. For each roughness level, we simulated three replicate landscapes (Appendix Figure 1). The generated landscapes were subsequently converted into binary habitat-matrix landscapes, considering different amounts of habitat *h* ranging from 0.05 to 0.4 in increments of 0.05, resulting in eight levels of habitat amount. Overall, we considered 120 distinct landscape configurations (5 roughness levels x 8 habitat amount levels x 3 replicates each). Each landscape was composed of 255 x 255 cells with cell resolution being adjusted to the spatial requirements of each species, ranging from 15 m to 35000 m spatial resolution, such that a high-quality habitat cell could support approximately 20 individuals.

### Dispersal of case study species

#### Dispersal in RangeShifter

RangeShifter simulates the three distinct phases of dispersal: emigration, transfer and settlement. In all case studies, both emigration and settlement are density-dependent, and in some even sex-dependent (Table 1, Appendix Figure 2). Settlement may require the presence of the opposite sex or is forced by reaching a maximal number of steps. All case studies used the stochastic movement simulator SMS to simulate the transfer process for each emigrating individual (Palmer et al. 2011). SMS assumes that moving through the landscape comes with specific costs (cost surface) for each habitat. These costs can be conceptualized as resistance against moving through a specific habitat rather than an actual cost to the individual. Individuals perceive the surrounding cost surface within their specific *perceptual range*. When moving through the landscape in single-cell steps without a predetermined destination, individuals base their decision to move to one of the neighbouring cells on the perceived surrounding cost surface, the *method* for evaluating the cost surface being case dependent, and their tendency to follow a correlated random walk. The correlated random walk is determined by the *directional persistence* and the *memory size*. With increasing directional persistence and memory size, individuals would keep moving in a certain direction, resulting in smoother movement paths. Additionally, a goal bias can be included so individuals will first move away from the natal location following a straighter path. As the distance from the natal location increases, as determined by the *dispersal bias decay rate* and *inflection point*, this tendency diminishes, resulting in less linear movement paths. Beyond movement costs, individuals may experience habitat specific step mortality, which is the probability of dying at each step of the transfer process.

**Figure 2:**
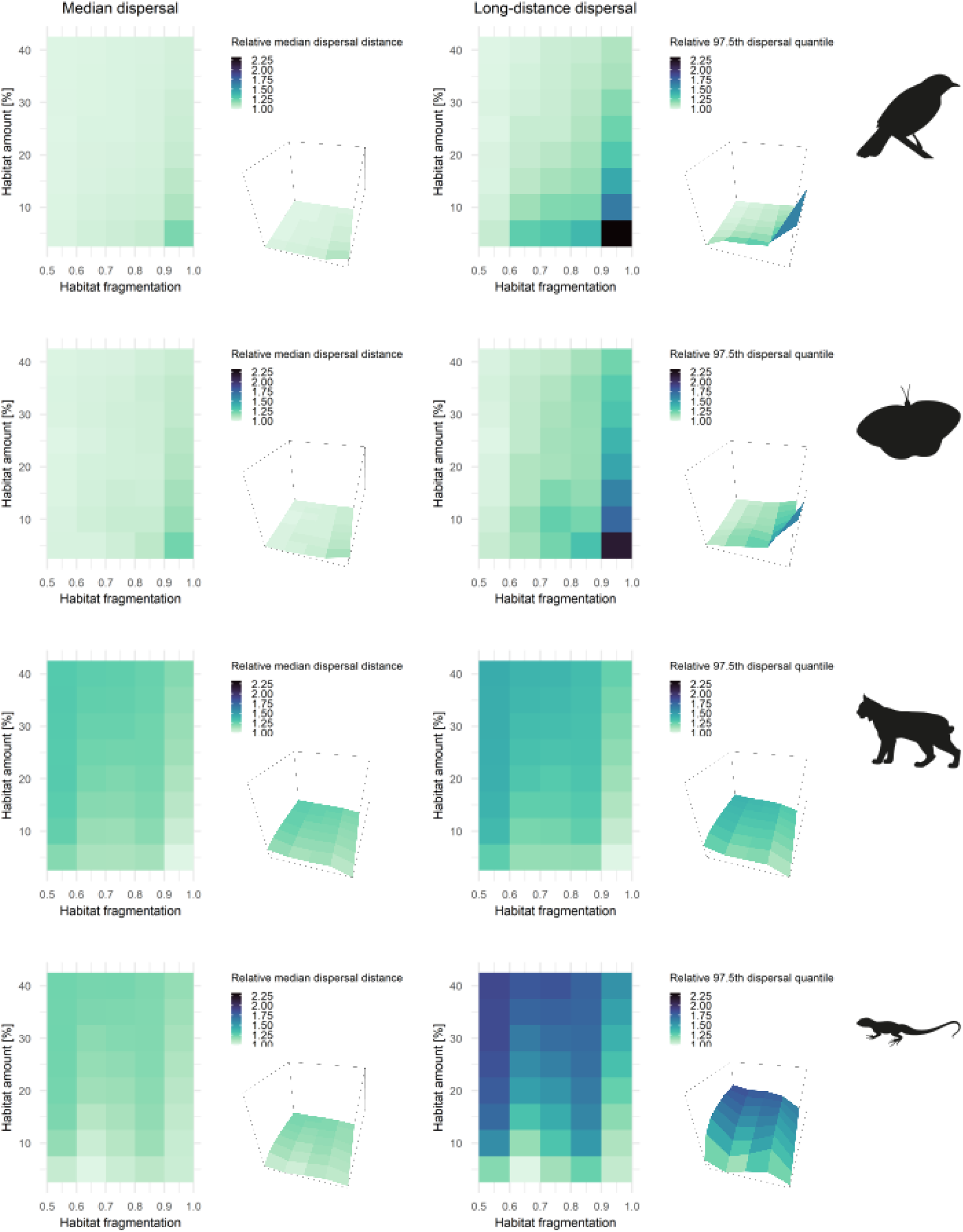
Median and long (97.5^th^ quantile) dispersal distances derived from bootstrapped samples of the fitted dispersal kernels for the four case study species (bird, insect, mammal and reptile) across different level of habitat amounts (in percentages) and degrees of fragmentation (expressed as roughness factor, r, with higher values indicating higher fragmentation, that is more small habitat patches). Dispersal distances are represented as mean distances over the three landscape replicates and are relative to the minimal median and long dispersal distance of each species. A value of 1 means equal to the minimal (median or long) dispersal distance, a value of 2 means doubling of the minimal (median or long) dispersal distance. These results are obtained using the log normal kernels for all species, that is the dispersal kernel which showed overall the highest weighted AIC value over all landscapes and species.

**Table 1:** Dispersal parameters for all 4 case study species. All RangeShifter parameters related to the dispersal process are listed. If dispersal processes are sex dependent, values for both sexes are given (f – female, m – male). Boolean parameters: T – true, F – false.

|  |  | Bird | Insect | Mammal | Reptile |
| --- | --- | --- | --- | --- | --- |
| Emigration | Probability | 0.7 (f), 0.5 (m) | 0.4 | 0.4 (f), 0.9 (m) | 0.8 (f), 0.9 (m) |
|  | Alpha | 0.6 (f), 0.6 (m) | 1 | 10 (f), 10 (m) | 10 (f), 10 (m) |
|  | Beta | 0.093 (f), 0.094 (m) | 5 | 1 (f), 1 (m) | 0 (f), 0 (m) |
|  | Sex dependency | T | F | T | T |
|  | Stage dependency | T | F | T | T |
|  | Density dependency | T | T | T | T |
| Transfer | Type | SMS | SMS | SMS | SMS |
|  | Perceptual range | 1 | 1 | 1 | 1 |
|  | Method | 2 | 2 | 2 | 2 |
|  | Memory size | 1 | 1 | 1 | 1 |
|  | Directional persistence | 2 | 5 | 5 | 5 |
|  | Goal bias type | 2 | 0 | 0 | 0 |
|  | Goal bias strength | 1.1 |  |  |  |
|  | Dispersal bias decay rate | 1 |  |  |  |
|  | Dispersal bias decay inflection point | 1000 |  |  |  |
|  | Costs [matrix, habitat] | 77, 1 | 100, 1 | 1000, 1 | 10, 1 |
|  | Per-step mortality probability | 1e-07 | 0.027 | 5e-04 (matrix), 0 (habitat) | 0.05 (matrix), 0 (habitat) |
|  | Straighten path | T | T | T | T |
| Settlement | Density dependency | T | T | T | T |
|  | Sex dependency | T | F | T | F |
|  | Stage dependency | F | F | F | F |
|  | Probability | 1 (f), 1 (m) | 1 | 1 (f), 1 (m) | 1 |
|  | Alpha (slope) | 100 (f), -150 (m) | -10 | -10 (f), -10 (m) | -10 |
|  | Beta (inflection point) | -1 (f), 0.055 (m) | 1 | 1 (f), 1 (m) | 0.7 |
|  | Minimal number of steps | 1 | 0 | 0 | 0 |
|  | Maximal number of steps | 12500 | 0 | 14 | 420 |
|  | Maximal number of steps per year | 0 | 0 | 0 | 40 |
|  | Mating requirements | T (f), F (m) | F | F (f), T (m) | F |

We based our work on previously published RangeShifter case studies of different animal species, including an insect (Melero et al. 2020), reptile (Williams et al. 2021), bird (Aben et al. 2016) and mammal (Ovenden et al. 2019), selected to encompass a range of dispersal abilities and population dynamics. The bird and insect species disperse volantly, but the bird species potentially disperses over longer distances and is less influenced by landscape features. Thus, having low resistance to fly over unfavourable habitat. Contrastingly, the reptile and mammal species, which disperse terrestrially, face higher costs and thus higher resistance when moving through unfavourable landscapes, e.g. urban areas, , with the mammal species showing a higher matrix resistance than the reptile species. Some of these case studies used patch-based landscapes, where habitat patches comprise clusters of cells with identical habitat. To avoid artefacts associated with dispersal in patch-based landscapes, where dispersal distance may be co-determined by how patches are defined and philopatric/allopatric behaviour, we reimplemented all case studies using a cell-based approach. Due to the switch to a cell-based model and the use of neutral habitat-matrix landscapes, we rescaled the spatial and habitat-dependent parameters cited in the original literature, particularly the perceptual range, memory size, movement costs, step mortality and maximal number of steps. A summary of the final dispersal settings is shown in Table 1, and a complete overview of all model parameters can be found in the appendix (Appendix 2: Parameter tables).

#### Bird case study

Aben et al. (2016) evaluated the effectiveness of conservation management scenarios for forest birds in a highly fragmented biodiversity hotspot. They found that increasing habitat area and connectivity through stepping stones can enhance population persistence and dispersal success, although spatial planning is essential to prevent creating dispersal sinks. We reimplemented their model using a cell-based approach, adjusting the cell resolution to 400 m to achieve stable population sizes of approximately 20 individuals per high-quality cell. Subsequently, we rescaled the spatial dispersal settings to align with the new cell resolution: perceptual range and memory size were each set to one cell and the maximal number of steps was set to 12500 cells. The cost value was kept at the highest value. Furthermore, the original model introduced dispersal bias in RangeShifter as a new feature but had not yet included all parameters defining dispersal bias of the current version RangeShifter 2.0 (Bocedi et al. 2021). Specifically, Aben et al. (2016) did not specify the dispersal bias decay rate and inflection point. To ensure the dispersal bias falls no lower than the value of the goal bias parameter, we set the decay rate to 1 and inflection point to a very high value (1000).

#### Insect case study

The insect case study is based on Melero et al. (2020), who utilized the RangeShifter modelling platform to assess the effectiveness of greening strategy in promoting urban biodiversity. They found that increasing the percentage of natural source areas around each garden was the most important factor for enhancing the colonization of butterfly species with different dispersal abilities. The original model was patch-based, but we adapted it to a cell-based approach, adjusting the cell resolution to 80 m. Consequently, the step mortality was multiplied by four (as the adjusted resolution is four times coarser, meaning one step represents 4 steps of the original resolution). The perceptual range and memory size were each set to one cell. Instead of permitting individuals to always settle when the habitat is suitable, we adjusted the settlement rule to include soft density dependence, with a probability of 1.0, slope of -10 and inflection point of 1. This modification prevents all individuals from immediately settling in a neighbouring habitat cell.

#### Mammal case study

In their study on reintroduction strategies for the Eurasian Lynx in Scotland, Ovenden et al. (2019) utilized the RangeShifter modelling platform to identify reintroduction sites with the highest probability of population persistence. We reimplemented the cell-based model with a cell resolution of 35 km. Accordingly, we rescaled the SMS parameters *perceptual range* and *memory size* to one cell each. The cost value for traversing matrix landscape was set to 1000. Furthermore, we adjusted the maximal number of steps to 14, thereby limiting individual movements to a maximum of 693 km.

#### Reptile case study

Williams et al. (2021) applied RangeShifter to simulate the range expansion of the invasive lizard species *Podarcis muralis* in England, UK. Due to climate suitability, the species’ range was restricted to the southern parts of the UK, while local landscape configuration and heterogeneity influenced population growth and range expansion. This suggested that regional expansion was likely to occur through exponential growth of local populations. As their RangeShifter implementation was already cell-based, we kept most of the original parameter values. However, as we used matrix-habitat landscapes assuming that matrix habitat was the most hostile habitat type, we set both the cost value for moving through the matrix and the step mortality in the matrix to the highest value specified in Williams et al. (2021), reflecting the most hostile habitat type.

### Simulations

All RangeShifter simulations were run for 50 years during which all case study species visually reached equilibrium (see Appendix Figure 3). For each of the 120 distinct landscapes (5 fragmentation levels x 8 habitat amount levels x 3 replicates each) we ran three RangeShifter replicate simulation runs. From the last year of each replicate run, we extracted start and end points of individual movement paths (of stage 0 for stage structured models and only successful emigrations), to ensure measuring natal dispersal for stable populations. These constitute observed dispersal distances and represent only natal dispersal. Uncertainty was captured from replicate landscapes and replicate RangeShifter simulations. We used the R package interface RangeShiftR (Malchow et al. 2021) for setting up and running all simulations.

**Figure 3:**
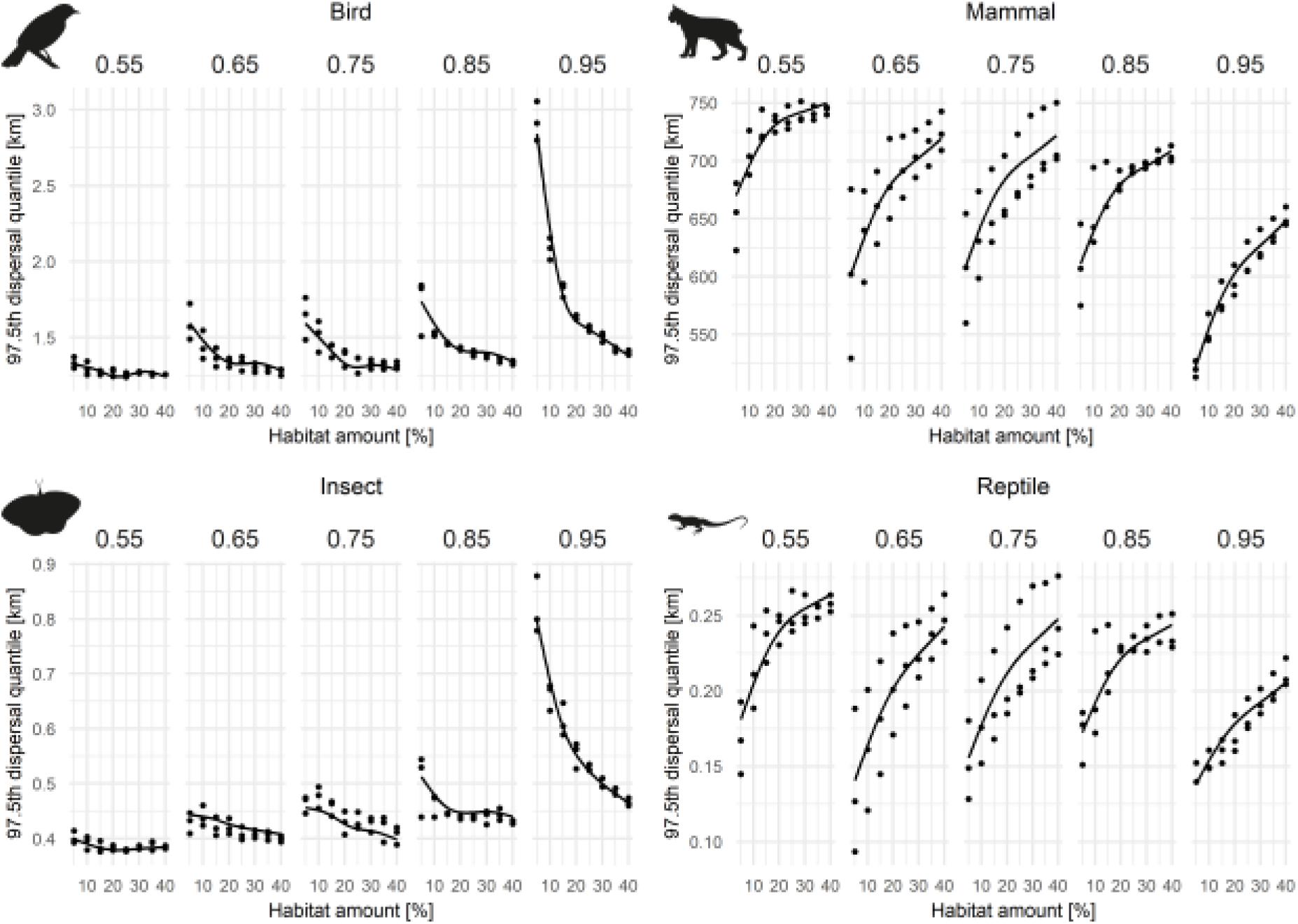
Impact of habitat amount (x-axis) and habitat fragmentation (columns) on the fitted (log normal) long dispersal distances (97.5^th^ quantile). Points show the fitted long dispersal distances for all landscape configurations and landscape replicates. Lines represent the fitted GAM model including habitat amount, habitat fragmentation and their interaction as predictors See Appendix Figure 6 and Appendix Figure 7 for the median and 99^th^ quantiles.

### Dispersal kernel estimation and context dependency

The simulated dispersal distances were subsequently used to fit common dispersal kernels for each species and landscape replicate. We applied maximum likelihood estimation to fit five dispersal kernel functions to the simulated dispersal distances of all successfully dispersed individuals (bird: N=14957-138219, insect: N=1651-15506, mammal: N=38467-376575, reptile: N=93707-802716) of the three replicates of each landscape configuration and repetition (using the fitdist function of the R package fitdistrplus, (Delignette-Muller and Dutang 2015)): Exponential, Weibull, Gamma, Half-Cauchy and Log-normal (Fandos et al. 2023). Mean weighted AIC values were used to identify the best-fitting distribution over all landscape configurations and repetitions. This was accomplished using the gofstat function of the R package fitdistrplus to calculate the AIC and the akaike.weights function of the R package qpcR (Ritz and Spiess 2008) to determine the weighted AIC. In all simulations – regardless of landscape replicate and species, log-normal distribution was the best fitting kernel. Thus, we proceeded in the subsequent analyses with the fitted log-normal kernels. From these kernels, we used bootstrapping to extract median and long-distance (97.5% and 99% quantiles) dispersal distances, thereby accounting for uncertainty in the estimated parameters.

To assess how dispersal quantiles vary across landscapes with different degrees of habitat fragmentation and habitat amount, we fitted generalized additive models (GAMs) using the gam function of the mgcv package (Wood 2003, 2004, 2011, 2017, Wood et al. 2016) with habitat amount, habitat fragmentation and their interaction as predictors. Abundance was not included due to high collinearity with habitat amount (correlation between habitat amount and abundance is larger than 0.99 for all species). We assessed the relative importance of each predictor using the gam.hp function of the R package gam.hp (Lai et al. 2024). The contribution of the interaction term (i.e. the joint contribution) was evenly distributed to each predictor. All analyses were conducted in R version 4.5.1 (2025-06-13 ucrt) (R Core Team 2025) using the packages NLMR for the generation of neutral landscapes (Sciaini et al. 2018), RangeShiftR for the RangeShifter simulations (Malchow et al. 2021), fitdistrplus for kernel estimations (Delignette-Muller and Dutang 2015), qpcR for weighted AIC calculations (Ritz and Spiess 2008), mgcv for the GAMs (Wood 2003, 2004, 2011, 2017, Wood et al. 2016), gam.hp for the relative importance of the predictors (Lai et al. 2024) and ggplot2 for figures (Wickham 2016). All codes are provided in a GitHub repository (see Data availability statement).

## Results

The simulated abundances of the four different species were strongly influenced by the amount of available habitat, with lower abundances in landscapes with reduced habitat amount (Appendix Figure 3). The reduction in abundance also led to fewer (successful) dispersers, driven by density-dependent processes during the dispersal (Appendix Figure 4). In contrast, varying degrees of habitat fragmentation had a relatively minor effect on species abundances and number of successful dispersers.

**Figure 4:**
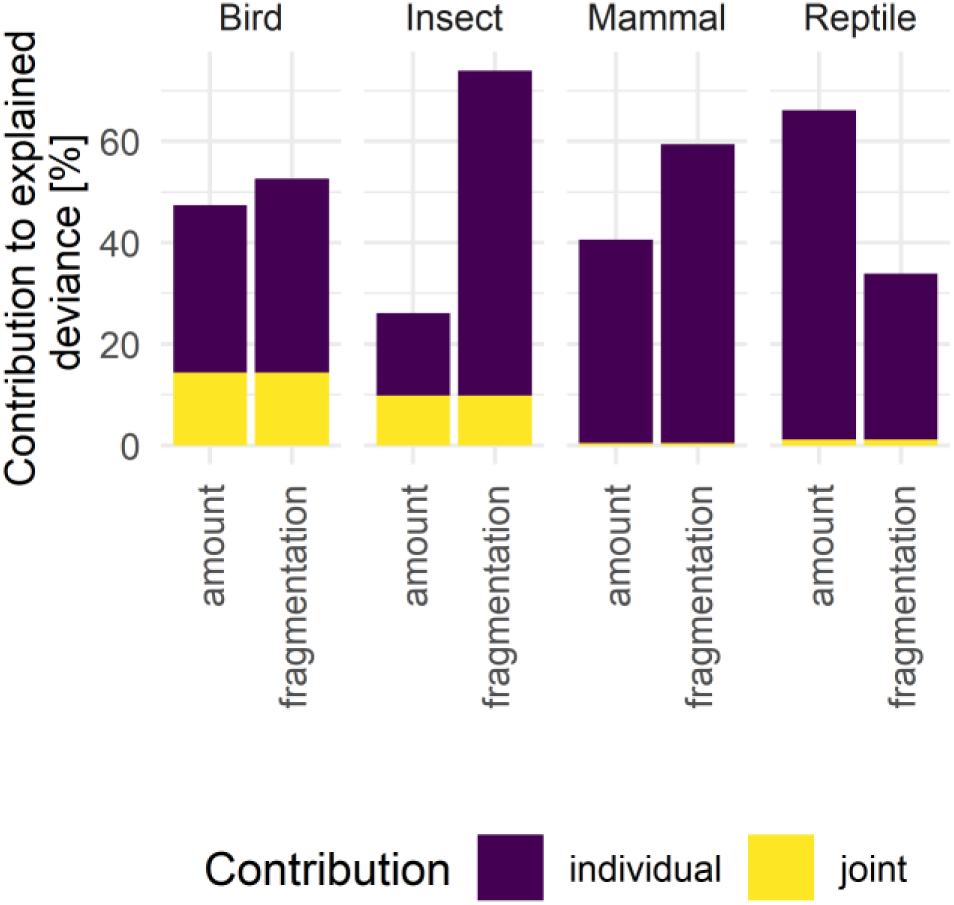
Variable importance of the GAM (97.5^th^ quantile ∼ s(amount) + s(fragmentation) + ti(amount, fragmentation)) based on the predictors’ contribution to the explained deviance. Abundance was excluded from the final model due to collinearity with habitat amount (correlation>0.99 for all species). Habitat fragmentation was the most important predictor for all case study species except for the reptile, for which habitat amount played the major role. The latter was also important for the bird and mammal species. For the mammal and reptile species, there was only a negligible joint contribution of both predictors. See Appendix Figure 8 and 9 for the median and 99^th^ quantiles, respectively.

The log-normal distribution was the best fitting kernel for all species and landscapes and we subsequently only report the results for this distribution. The effect directions of landscape configuration were similar for the median and long (97.5^th^ quantile) dispersal distances (Figure 2, Appendix Figure 5). For the bird and insect case study species, longer distances emerged in landscapes with high fragmentation and low habitat amount. This pattern was even more pronounced for long-distance dispersal compared to median dispersal distance. However, the mammal and reptile case study revealed an opposite pattern with dispersal distances decreasing in more fragmented landscapes with lower habitat amount. A generally negative relationship between dispersal distance and habitat amount was observed for the bird and insect case studies, whereas a generally positive relationship was found for the mammal and reptile case study (Figure 3).

For both the bird and insect, the negative relationship between dispersal distance and habitat amount intensified with increasing habitat fragmentation, showing a strong non-linear, negative response at the highest degree of habitat fragmentation, while no relationship was evident at the lowest degree of fragmentation (Figure 3). In contrast, the reptile and mammal case study species showed a consistently positive relationship between dispersal distance and habitat amount, regardless of the degree of habitat fragmentation. However, this relationship became steeper and more linear as habitat fragmentation increased.

For most case studies, habitat fragmentation was the predominant factor explaining differences in long-distance dispersal patterns between landscapes (Figure 4), especially for the insect case study, where habitat fragmentation accounted for more than 70% of the explained deviance. In two out of the four species, the interaction between habitat amount and habitat fragmentation, represented by their joint contribution, played an important role in shaping dispersal patterns highlighting the complex interplay of habitat components (bird and insect). For the mammal and reptile case study species, the interaction was negligible. The reptile was the only case study species for which habitat amount was the dominant factor explaining the difference in dispersal distances. These patterns were consistent across different dispersal distances (median, 97.5^th,^ and 99^th^ quantiles; Appendix Figure 6-9).

## Discussion

Our study provides a deeper theoretical understanding of the context dependency of dispersal. It demonstrates that dispersal kernels are not fixed species traits but influenced by individual movement behaviour and landscape configuration. Across four taxonomically distinct species, we consistently found log-normal distributions (characterized by fat tails in our parameterization), indicating a persistent signal of rare long-distance dispersal. Yet, the scale and shape of dispersal were strongly contingent on habitat amount and fragmentation, with fragmentation generally having a stronger effect on emergent dispersal distances than habitat amount. Importantly, species differed in their responses to these landscape factors, reflecting differences in vagility and matrix resistance. These findings highlight that intraspecific variation in dispersal distances can be expected across landscapes, emphasizing the need to treat dispersal as context dependent rather than transferable across environments.

Our findings contribute to the long-standing debate over the relative roles of habitat loss and fragmentation per se (Fahrig 2003; Haddad et al. 2015). In line with recent syntheses (Valente et al. 2023; Sgarlata et al. 2025), we found that habitat loss consistently reduced abundances across taxa, whereas fragmentation had relatively little effect on population size but strongly influenced emergent dispersal kernels. This distinction underscores that while habitat loss drives demographic decline, fragmentation per se alters the spatial scale and shape of dispersal, with consequences for connectivity and range dynamics. Importantly, the impact of fragmentation was not uniform but depended on species’ vagility and matrix resistance, highlighting the context-dependence emphasized in recent contributions to the debate (Riva 2025). By mechanistically linking dispersal behaviour to landscape structure, our study provides a process-based perspective on when and how fragmentation effects are likely to matter, offering a bridge between pattern-focused landscape metrics and predictions of population responses under global change.

Across most simulated taxa, habitat fragmentation was the main determinant of the observed context dependency in dispersal (Figure 4). As hypothesized, high-vagility species (birds, insects) tend to increase dispersal with fragmentation, whereas low-vagility species (mammals and reptiles) tend to decrease dispersal. These species-specific responses of dispersal distance to landscape configuration can be explained by species’ perception of the landscape, the associated constraints and the mortality risk during dispersal. Highly vagile species such as birds and insects experience the landscape matrix as relatively permeable (easy to pass through), with mortality risks being less dependent on the habitat, and thus are less constrained to cross low-quality habitats when patch distances increase. For example, nuthatches showed greater mean dispersal distances in heterogenous landscapes compared to dense forests (Matthysen et al. 1995). In contrast, non-volant and low-vagility species, such as the reptile and mammal, perceive higher risks and often avoid crossing unfavourable habitat due to increased movement costs or higher mortality risks, e.g. in urban areas. Our mammal case study species, the Eurasian lynx, for example, strongly avoids urban habitats where mortality risk is higher due to road kills (Zimmermann and Breitenmoser 2007). Increasing edge to area ratios in fragmented landscapes confronts dispersing individuals more frequently with habitats of high movement costs or mortality risks and thereby restricting individual movements in general, especially for mammals, where edges can actually represent physical semi-permeable barriers like roads or fences potentially restricting the movement (Robb et al. 2022).

In our models, the species-specific risk perception was simplified to a fixed (species-specific) cost value and step mortality risks for the mammal and reptile case study species. However, in reality, risk perception is a complex process and both risk perception and risk mortality depend on species-specific traits (dispersal mode, body size and mobility, defence mechanisms, physiology or individual behaviour (Stevens et al. 2014)) as well as landscape features (e.g. barriers, resources and potential shelter). White-tailed deer, for example, dispersed farther in more open habitat compared to forested areas, showing a trade-off between finding cover in forests and detecting predators in open habitats (Long et al. 2005), both influencing the overall risk perception of the habitat. Our findings underscore that dispersal is an emergent property of individual movement decisions and risk perception interacting with landscape structure, with important consequences for modelling connectivity and population dynamics. Recent studies show that beyond risk perception, individuals’ survival in the matrix is a key determinant of landscape connectivity (Day et al. 2020, Yamaura et al. 2022, Argote-Deluque et al. 2025). Landscape-mediated variability in dispersal across populations corroborates previous findings (Schtickzelle et al. 2006, Jacob et al. 2020) hampering the transferability of dispersal kernels to new landscape contexts or populations. This can introduce significant uncertainties in spatial population modelling and predictions of population spread, for example under global change, when habitat conditions change over time, or following conservation and restoration measures. Mechanistic movement models, such as the stochastic movement model used in this study, can partially address these uncertainties by simulating movement decisions based on the perception of landscape features, thereby capturing context-dependent behaviour and the emerging dispersal differences across populations and landscapes (Aben et al. 2016, Allen et al. 2016, Hunter-Ayad and Hassall 2020). Yet, intraspecific variability due to adaptation or evolution in movement traits could further exacerbate context dependency but would be more difficult to anticipate and parameterize. Future research could focus on studying how evolution of movement-related traits and intra-specific variability in these traits influence the reported relationships between dispersal distance and habitat fragmentation and amount.

Despite context-dependent differences in dispersal, our results highlighted a consistent role of long-distance dispersal (LDD) events. Also empirically, LDD was observed for plants (Austerlitz et al. 2004), birds (Van Houtan et al. 2007, Kesler et al. 2010, Fandos et al. 2023) and beetles (combining empirical data with modelling, Chapman et al. 2007). However, especially in the field, LDD might be underestimated. If individuals leave a study area in a mark-recapture experiment or tracking devices are lost, it remains unknown whether this is because individuals died or dispersed beyond the study area. Detecting LDD empirically is even more tricky as only up to 5% of emigrating individuals are expected to exhibit LDD and sample sizes in mark recapture or tracking studies is in many cases too small for robust estimates (Chadœuf et al. 2017, Fandos et al. 2023). On the other hand, GPS tracking data provide valuable insights into individual decisions during the transfer phase and are thus particularly valuable for parameterizing stochastic movement simulators as used in RangeShifter allowing to simulate emerging dispersal distances rather than relying on dispersal kernels (Palmer et al. 2011). Unfortunately, current knowledge on species movements across taxonomic groups is limited (Scarpignato et al. 2023) and large data sets on empirical dispersal kernels or movement data remain scarce (Bullock et al. 2017, Chu and Claramunt 2023, Fandos et al. 2023). We suggest that both types of data, mark-recapture data across many individuals and high-resolution tracking data across fewer individuals, are essential for understanding dispersal and its context dependency, and for inclusion in ecological models of spatial population dynamics.

Our findings underscore a fundamental challenge for predicting species responses to environmental change. Dispersal kernels are emergent properties of species–landscape systems rather than fixed species traits. The magnitude of context-dependency we observed, particularly for low-vagility species in fragmented landscapes, suggests that transferring kernels across environments risks substantial prediction error. Mechanistic movement models offer a promising path forward by explicitly linking individual behaviour to landscape structure (Zurell et al. 2022), but their power ultimately depends on integrating empirical data. High-resolution tracking can parameterize movement rules, and population-level mark–recapture can validate emergent patterns. Bridging theoretical models with these complementary empirical approaches will be essential for building predictive frameworks that can anticipate species persistence in increasingly human-modified landscapes.

## Data availability statement

We provide a git repository with all codes to reproduce the simulations, analyses and figures: https://anonymous.4open.science/r/XXXX_dispersal_kernel_2022-B39E

## Conflict of interest

The authors declare that they have no conflict of interest.

## Author contributions

Conceptualization: JW, DZ

Analysis: JW

Methodology: JW, DZ, GF

Writing – original draft preparation: JW, DZ

Writing – review & editing: all authors

## Notes

### Competing Interest Statement

The authors have declared no competing interest.

