## Appendix for "Landscape context drives within-species variation in dispersal kernels and limits transferability"

### Supplementary material

#### Appendix 1: additional figures and tables

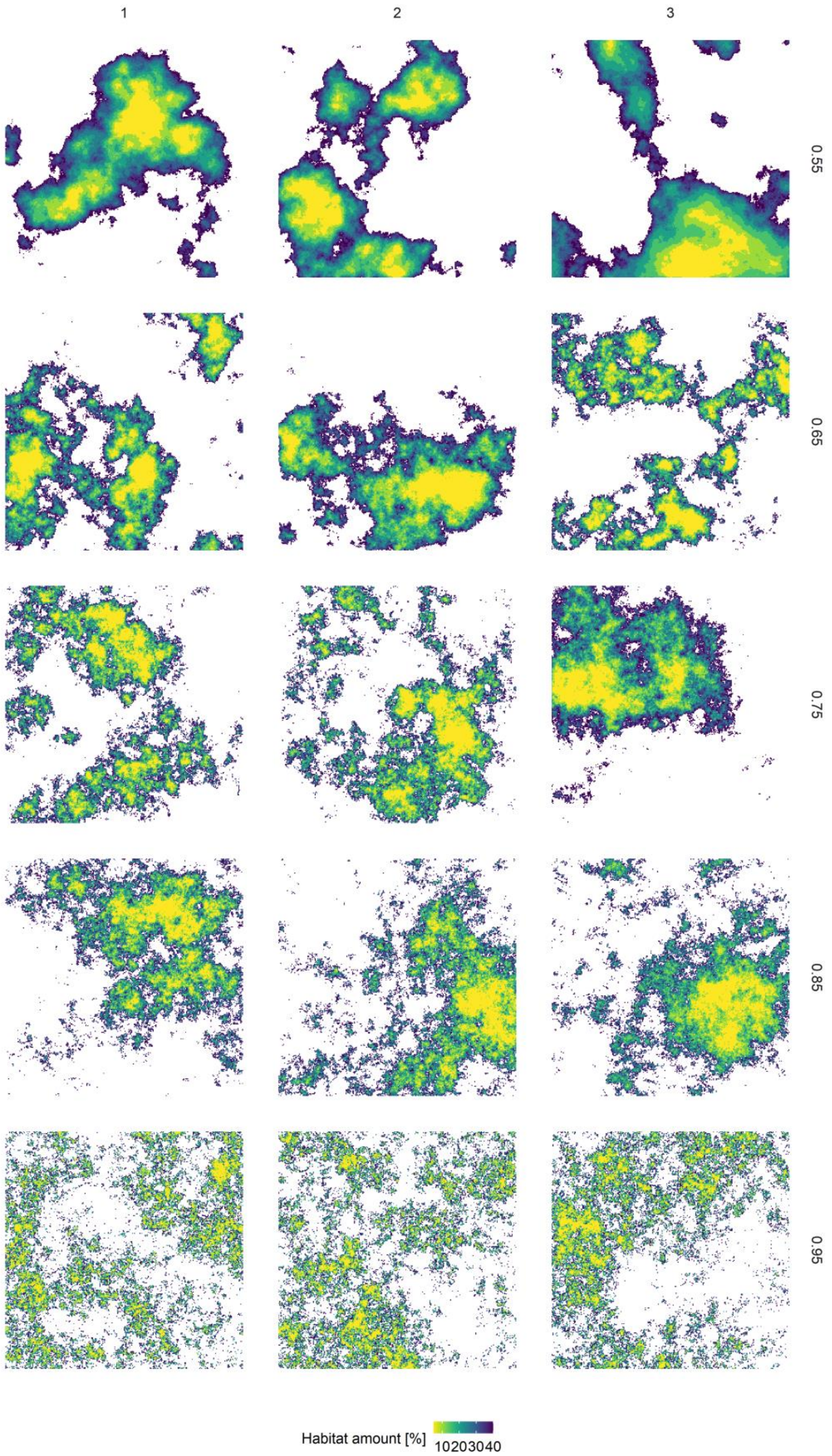

Appendix Figure 1: Generated neutral landscapes with different habitat fragmentations (5 levels, rows) and habitat amounts (8 levels, colour). Landscape replicates are shown in columns. Fragmentation, the degree of spatial autocorrelation, can take values between 0 and 1; a value of 1 corresponding to the lowest autocorrelation indicating highest fragmentation with many small habitat patches.

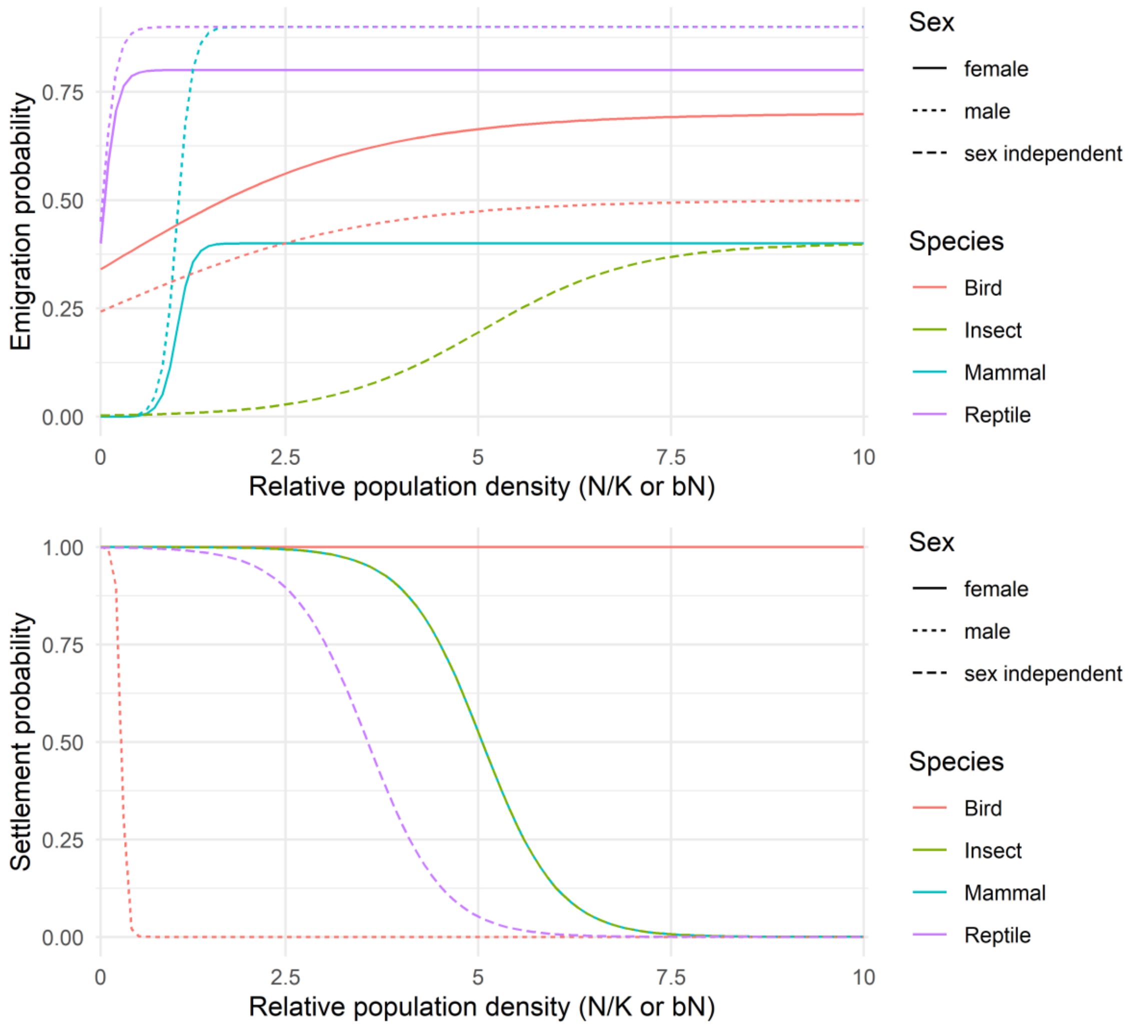

Appendix Figure 2: The species specific effect of density dependence on emigration and settlement probabilities.

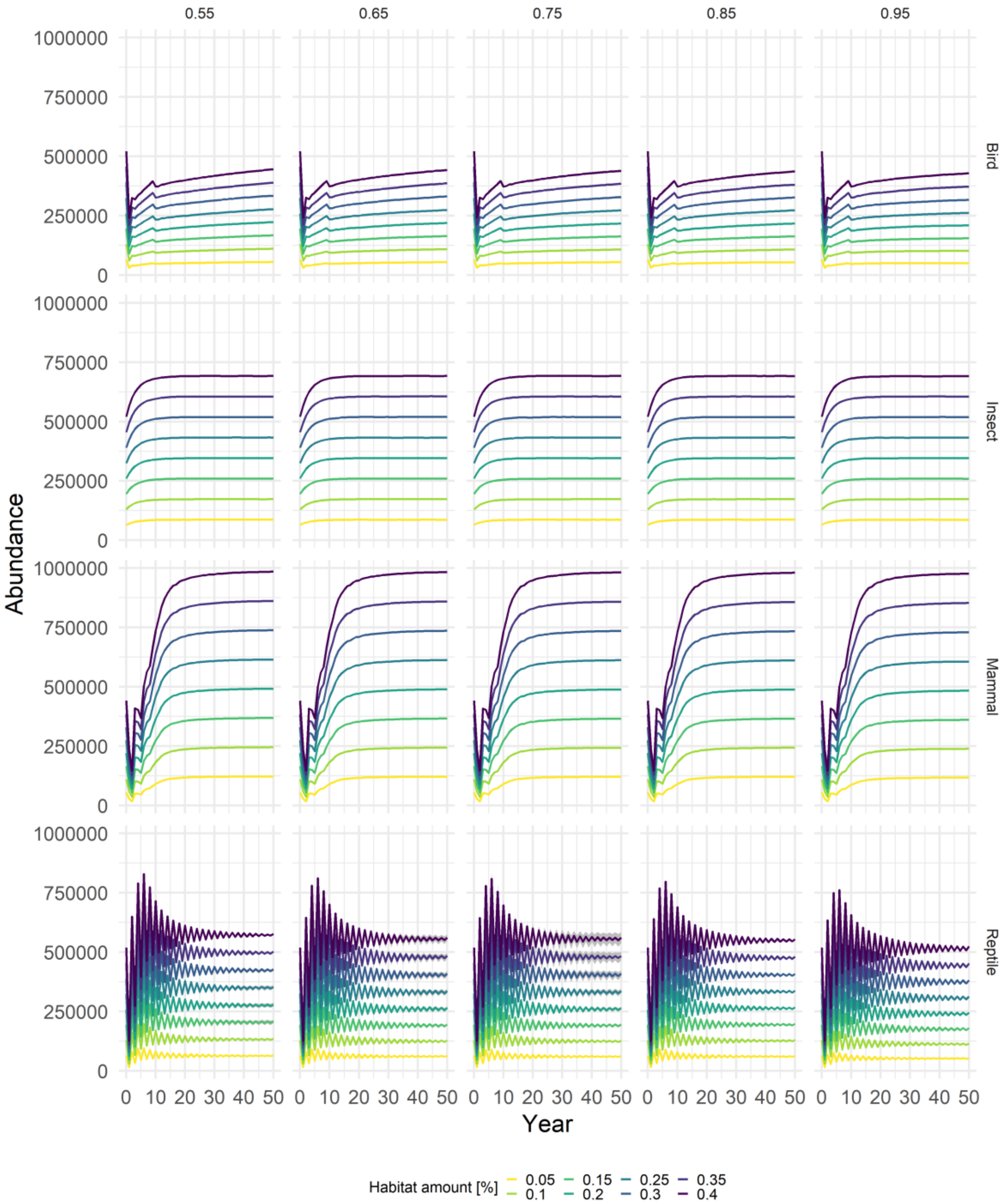

Appendix Figure 3: Abundances of the four case study species in different landscape configurations. Lines show the mean abundances over three RangeShifter replications and three landscape repetitions, ribbons show the standard deviation. Colours code for habitat amount, rows show the different habitat fragmentations with higher values indicating higher fragmentation (i.e. many small patches, which are more evenly distributed). Please note that for fitting dispersal kernels we used the sum of successful dispersers over the three RangeShifter replicates.

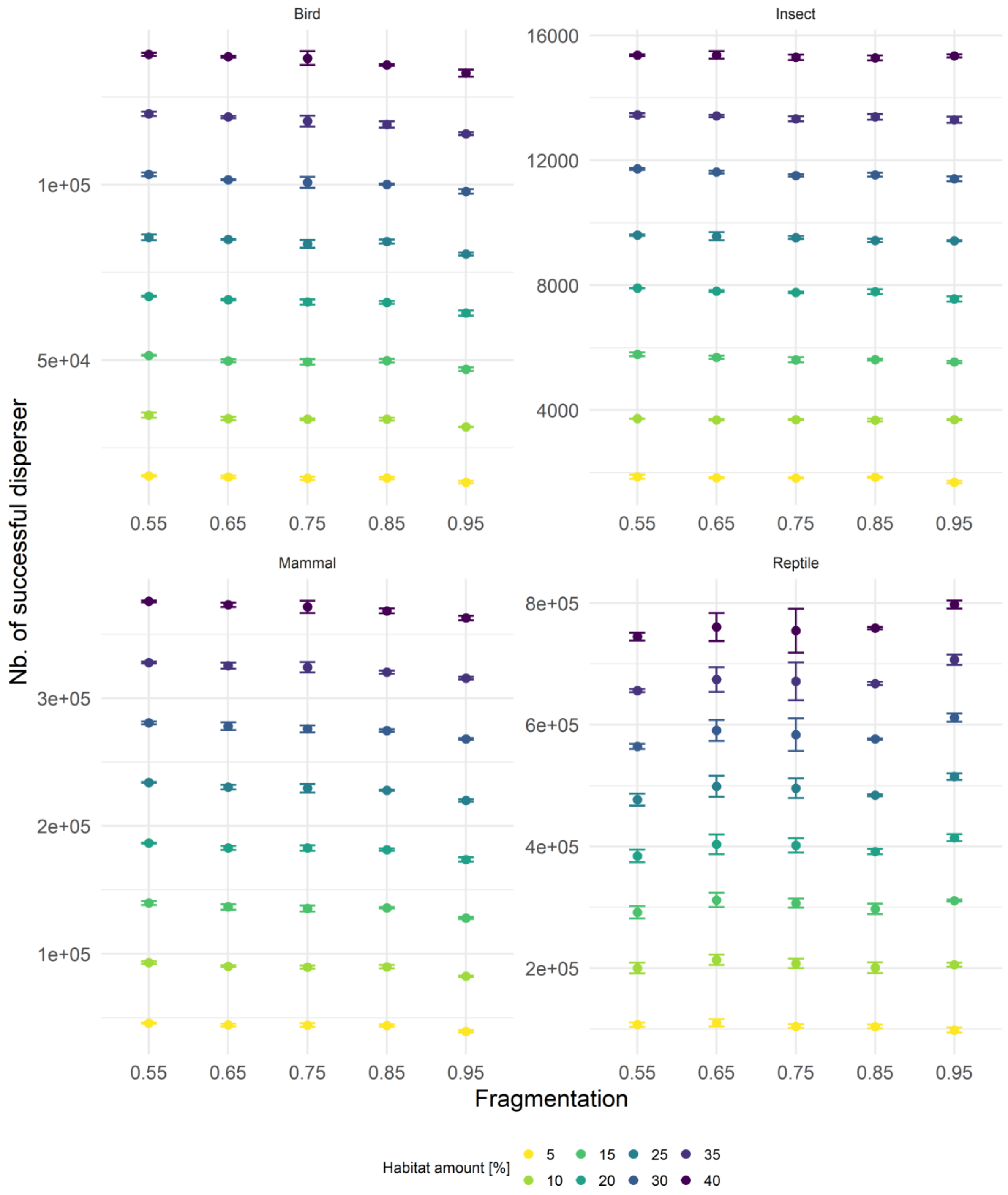

Appendix Figure 4: Number of successful disperser of the four case study species in different landscape configurations. Points show the mean number of dispersers over three landscape repetitions, error bars show the standard deviation. Note that we used the sum of all successful disperser in three RanggeShifter replicates. Colours code for habitat amount and the x axis shows the different habitat fragmentations with higher values indicating higher fragmentation (i.e. many small habitat patches, which are more evenly distributed in the landscape.

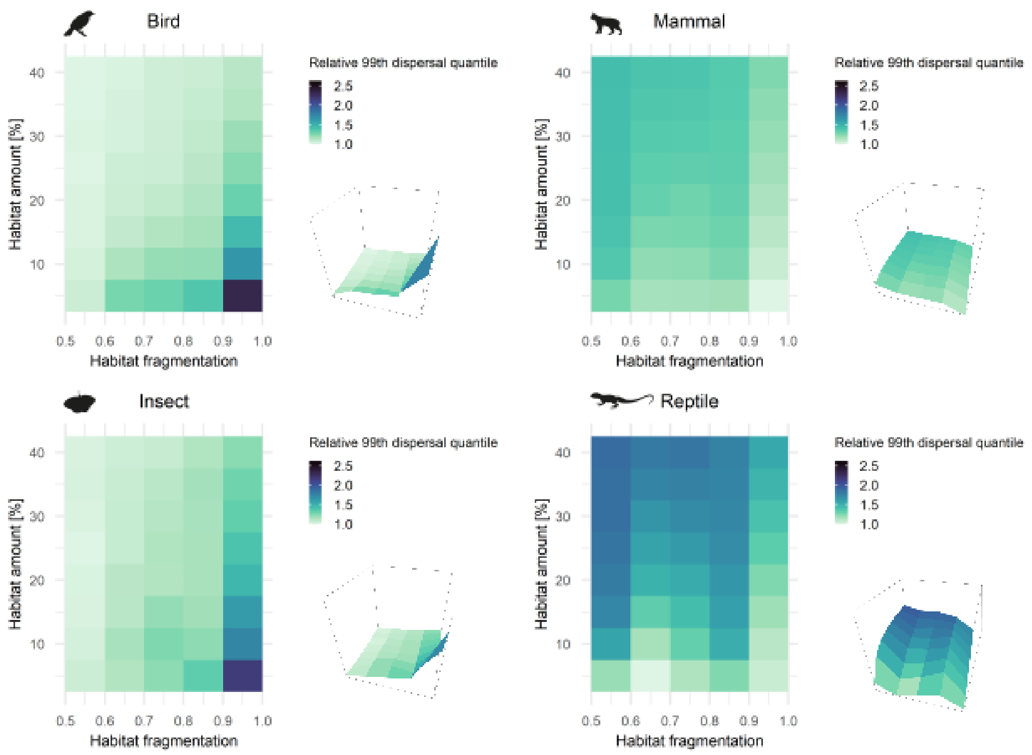

Appendix Figure 5: Long dispersal distances (99$^{th}$ quantile) of the four case study species. Colours show the relative 99$^{th}$ quantile of bootstrapped samples from the fitted log-normal kernel. Log-normal kernel was used as it showed in all case studies the highest weighted AIC. Relative values were calculated by dividing the species-specific quantiles by the minimum value of the specific species.

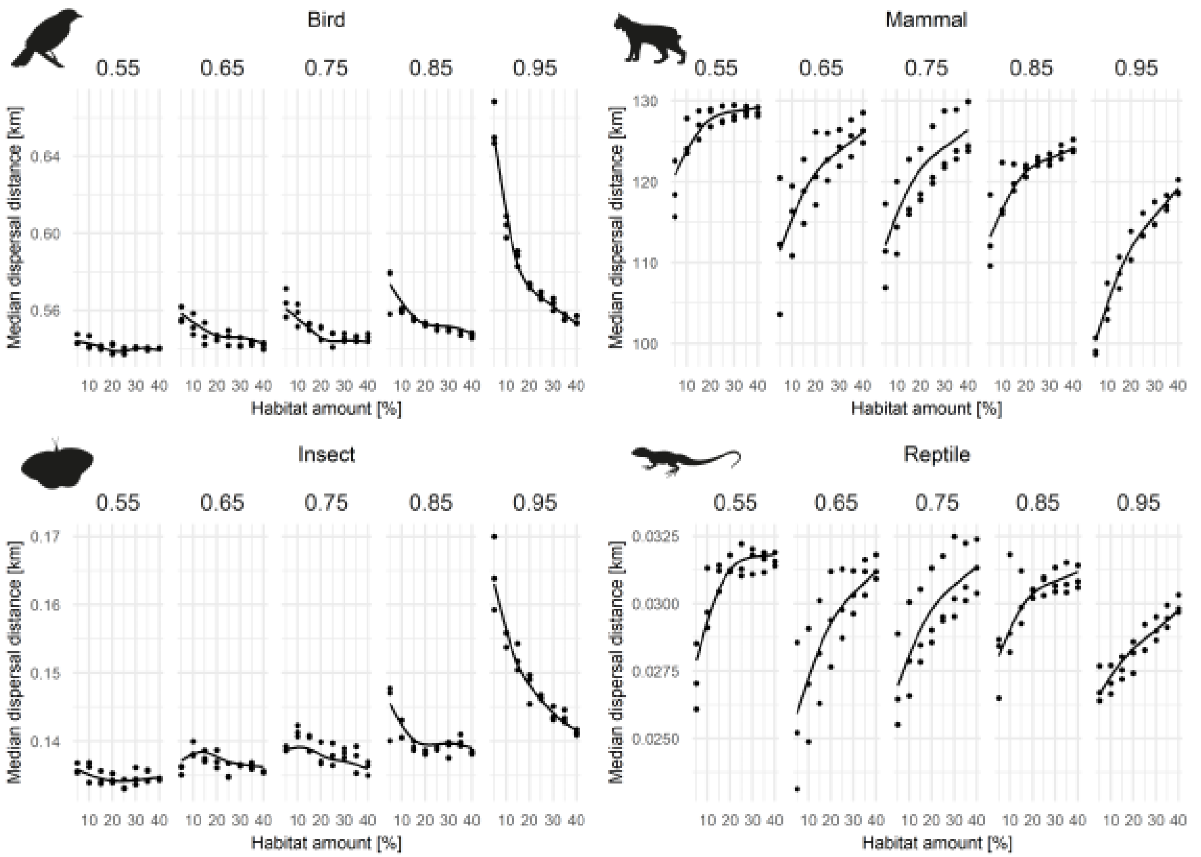

Appendix Figure 6: Impact of habitat amount and habitat fragmentation on the estimated median dispersal quantiles obtained from bootstrapped samples from the fitted log-normal distributions. Points show the estimated median dispersal distances for all landscape configurations and landscape replicates. Lines represent the final model including habitat amount, habitat fragmentation and interaction between amount and fragmentation (gam($50^{th}$ quantile ~ s(amount) + s(fragmentation) + ti(amount, fragmentation)). We did not include abundance or number of dispersers as both were highly correlated with habitat amount (correlation > 0.99).

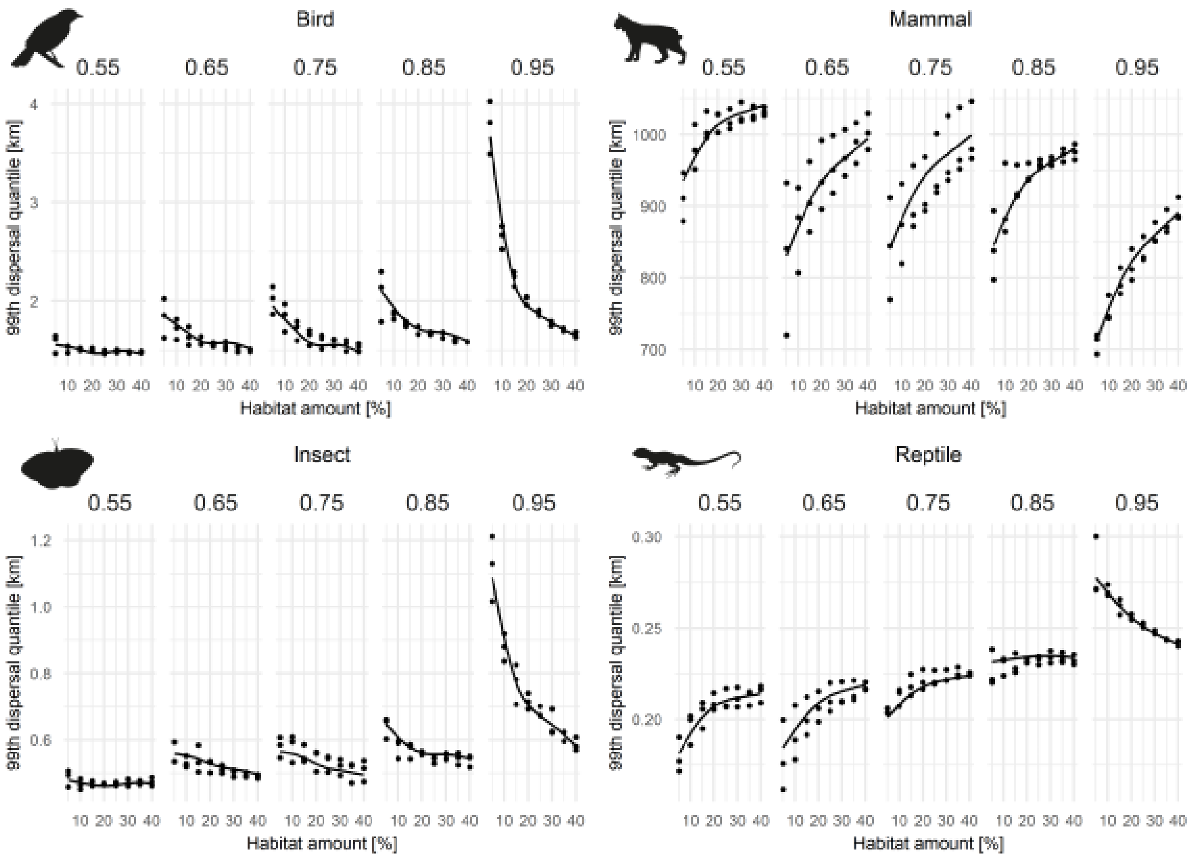

Appendix Figure 7: Impact of habitat amount and habitat fragmentation on the estimated 99th dispersal quantiles obtained from bootstrapped samples of the fitted log-normal distribution. Points show the estimated 99th dispersal quantiles for all landscape configurations and landscape replicates. Lines represent the final model including habitat amount, habitat fragmentation, and interaction between amount and fragmentation (gam(99th quantile ~ s(amount) + s(fragmentation) + ti(amount, fragmentation)). We did not include abundance or number of dispersers as both were highly correlated with habitat amount (correlation > 0.99).

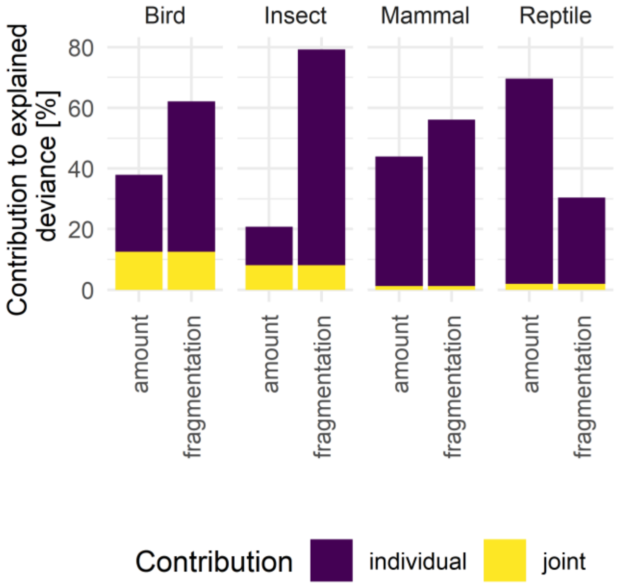

Appendix Figure 8: Variable importance of the statistical model of the median dispersal distances (gam(50th quantile ~ amount + fragmentation + ti(amount, fragmentation)) for all case study species. Variable importance was caclucated using the gam.hp function of the R package gam.hp (Lai et al. 2024). The contribution of the interaction term was evenly distributed to each predictor. Habitat fragmentation was in all cases but the reptile the most important factor. Contrastingly, habitat amount was more important for the reptile case study species.

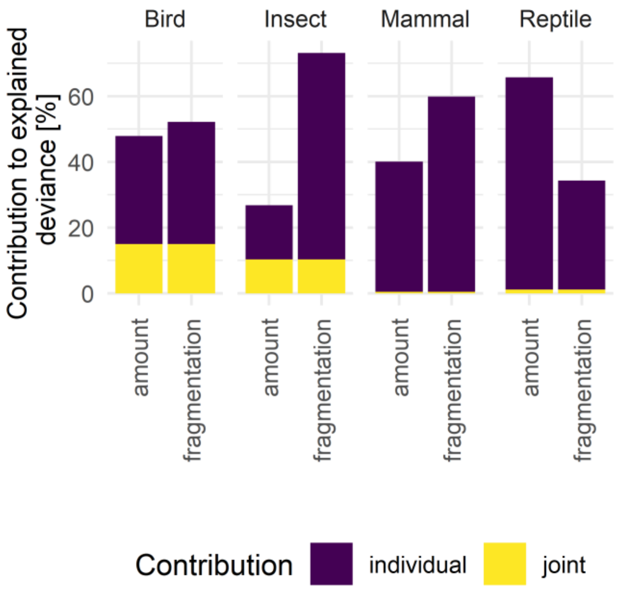

Appendix Figure 9: Variable importance of the statistical model (gam(99$^{th}$ quantile ~ s(amount) + s(fragmentation) + ti(amount, fragmentation)) for all case study species. Variable importance was calculated using the gam.hp function of the R package gam.hp (Lai et al. 2024). The contribution of the interaction term was evenly distributed to each predictor. Habitat fragmentation was in all cases but the reptile the most important factor. Contrastingly, habitat amount was more important for the reptile case study species.

Appendix Table 1: Mean weighted AIC values for the five fitted dispersal kernels. Mean was calculated over all landscape configurations and replicates. In all cases, log-normal distribution showed by far the best fit. AIC difference was so high that the weighted AIC was always 1.

| Species | Kernel | Weighted AIC | |
|---|---|---|---|
| | | Mean | SD |
| Mammal | Weibull | 0 | 0 |
| | Gamma | 0 | 0 |
| | Log normal | **1** | 0 |
| | Half-cauchy | 0 | 0 |
| | Exponential | 0 | 0 |
| Bird | Weibull | 0 | 0 |
| | Gamma | 0 | 0 |
| | Log normal | **1** | 0 |
| | Half-cauchy | 0 | 0 |
| | Exponential | 0 | 0 |
| Reptile | Weibull | 0 | 0 |
| | Gamma | 0 | 0 |
| | Log normal | **1** | 0 |
| | Half-cauchy | 0 | 0 |
| | Exponential | 0 | 0 |

| | | | |
|---|---|---|---|
| Insect | Weibull | 0 | 0 |
| | Gamma | 0 | 0 |
| | Log normal | **1** | 0 |
| | Half-cauchy | 0 | 0 |
| | Exponential | 0 | 0 |

### Appendix 2: Parameter tables

#### Bird case study

##### Simulation parameters

| Parameter | Description | Values |
|---|---|---|
| Year | Number of simulated years | 50 |
| Replicates | Number of simulation iterations | 3 |
| Absorbing | Whether non-valid cells lead to direct mortality of the individual during transfer | FALSE |
| LocalExt | Local extinction | FALSE |
| EnvStoch | Environmental stochasticity (0: none, 1: global, 2: local) stochasticity acts (0: growth rate/fecundity, 1: demographic density dependence) | 0 |
| OutIntRange | Output of range file | 1 |
| OutIntOcc | Output of occupancy file | 0 |
| OutIntPop | Output of population file | 0 |
| OutIntInd | Output of individual file | 1 |
| OutIntConn | Output of connectivity file | 0 |
| OutIntPaths | Output of SMS paths file | 0 |
| OutStartPop | Starting year for output population file | 0 |
| OutStartInd | Starting year for output individual file | 49 |
| OutStartConn | Starting year for output connectivity file | 0 |
| OutStartPaths | Starting year for output SMS paths file | 0 |
| SMSHeatMap | Output SMS heat map raster file | FALSE |

##### Landscape parameters

| Parameter | Description | Values |
|---|---|---|
| Landscape file | Filename(s) of the landscape map(s) | |
| Resolution | Resolution in meters | 400 |
| HabPercent | Whether habitat types/codes or habitat cover/quality | FALSE |
| NHabitats | Number of different habitat codes | 2 |
| K_or_DensDep | Demographic density dependence | 0, 28.986 |
| PatchFile | Filename(s) of the patch map(s) | |
| CostsFile | Filename(s) of the SMS cost map(s) | |
| DynamicLandYears | Years of landscape changes | 0 |
| SpDistFile | Filename of the species initial distribution map | |

Demography parameters

| Parameter | Description | Values |
|---|---|---|
| Stages | Number of life stages | 3 |
| TransMatrix | Transition matrix. Defines the development probabilities from each stage into the next as well as the respective survival probabilities and fecundities | $\begin{bmatrix} 0 & 0 & 0 & 0 & 1 & 1.38 \\ 1 & 0 & 0 & 0 & 0 & 0 \\ 0 & 1 & 0 & 0 & 0 & 0 \\ 0 & 0 & 0.508 & 0 & 0.85 & 0 \\ 0 & 0 & 0 & 0.463 & 0 & 0.825 \end{bmatrix}$ |
| MaxAge | Maximum age in years | 10 |
| MinAge | Ages which an individual in stage i-1 must already have reached before it can develop into the next stage i. | 0, 0, 0, 0, 0 |
| RepSeasons | Number of potential reproduction events per year | 1 |

| Parameter | Description | Values |
|---|---|---|
| RepInterval | Number of reproductive seasons which must be missed following a reproduction attempt before another reproduction attempt may occur | 0 |
| PRep | Probability of reproducing in subsequent reproductive seasons | 1 |
| SurvSched | Scheduling of survival (0: at reproduction, 1: between reproductive events, 2: annually) | 1 |
| FecDensDep | whether density dependent fecundity probability is modelled | FALSE |
| DevDensDep | Whether density dependent development probability is modelled | FALSE |
| SurvDensDep | Whether density dependent survival probability is modelled | TRUE |
| DevDensCoeff | Relative density dependence coefficient for development | 1 |
| SurvDensCoeff | Relative density dependence coefficient for survival | 1 |
| FecStageWtsMatrix | Stage-dependent weights in density dependence of fecundity | Not selected. |
| DevStageWtsMatrix | Stage-dependent weights in density dependence of development | Not selected. |
| SurvStageWtsMatrix | Stage dependent weights in density dependence of survival | Not selected. |

| Parameter | Description | Values |
|---|---|---|
| PostDestrictn | Whether individuals of a population die (FALSE) or disperse (TRUE) if its patch gets destroyed | FALSE |
| ReproductionType | Decribes the reproduction type (0: asexual/only female; 1: simple sexual model; 2: sexual model with explicit mating system) | 2 |
| PropMales | Proportion of males in the population | 0.5 |
| Harem | Maximum harem size | 1 |

Dispersal parameters

| Process | Parameter | Description | Values |
|---|---|---|---|
| Emigration | EmigProb | Emigration probabilities/parameters for each stage/sex | $\begin{bmatrix} 0 & 0 & 0.7 & 0.6 & 0.093 \\ 0 & 1 & 0.5 & 0.6 & 0.094 \\ 1 & 0 & 0 & 0 & 0 \\ 1 & 1 & 0 & 0 & 0 \\ 2 & 0 & 0 & 0 & 0 \\ 2 & 1 & 0 & 0 & 0 \end{bmatrix}$ |
| | SexDep | Sex-dependent emigration probability? | TRUE |
| | StageDep | Stage-dependent emigration probability? | TRUE |
| | DensDep | Density-dependent emigration probability? | TRUE |
| | UseFullKern | Shall the emigration probability be derived from dispersal kernel? | FALSE |
| Transfer | Type | Type of transfer | StochMove |
| | PR | Perceptual range | 1 |
| | PRMethod | Method to evaluate the effective cost of a particular step from the landdscape | 2 |

| Process | Parameter | Description | Values |
|---|---|---|---|
| | | within the perceptual range:<br>1: Arithmetic mean<br>2: Harmonic mean<br>Weighted arithmetic mean | |
| | MemSize | Size of memory, given as the number of previous steps<br>over which to calculate current direction<br>to apply directional persistence | 1 |
| | DP | Directional persistence. Corresponds to the<br>tendency to follow a correlated random walk. | 2 |
| | GoalType | Goal bias type<br>0: None, 2: dispersal bias | 2 |
| | GoalBias | Goal bias strength | 1.1 |
| | AlphaDB | Dispersal bias decay rate | 1 |
| | BetaDB | Dispersal bias decay inflection point (number of steps) | 1000 |
| | Costs | Describes the landscapes resistance to movement:<br>Either habitat-specific costs for each habitat type<br>or 'file', to indictae to use the cost raster map(s)<br>specified in the landscape module. | 77, 1 |
| | StepMort | Per-step mortality probability: constant or habitat-specific | 1e-07 |
| | StraightenPath | Straighten path after decision not to settle in a patch? | TRUE |

| Process | Parameter | Description | Values |
|---|---|---|---|
| Settlement | DensDep | Density-dependent settlement requirements? | TRUE |
| | SexDep | Sex-dependent settlement requirements? | TRUE |
| | StageDep | Stage-dependent settlement requirements? | FALSE |
| | Settle | Settlement probability parameters for all stages/sexes. | $\begin{bmatrix} 0 & 1 & 100 & -1 \\ 1 & 1 & -150 & 0.055 \end{bmatrix}$ |
| | MinSteps | Minimum number of steps | 1 |
| | MaxSteps | Maximum number of steps | 12500 |
| | MaxStepsYear | Maximum number of steps per year | 0 |
| | FindMate | Mating requirements to settle? | TRUE, FALSE |

#### Initialisation parameters

| Parameter | Description | Values |
|---|---|---|
| InitType | Type of initialisation<br>(0: free initialisation according to habitat map,<br>1: from loaded species distribution map,<br>2: from initial individuals list file) | 0 |
| FreeType | Option for free initialisation<br>(0: random in given number of cells,<br>1: all suitable cells/patches)<br>species distribution map (0: all suitable cells within<br>all distribution presence cells,<br>1: all suitable cells within given<br>number of randomly chosen presence cells) | 1 |
| InitDens | Number of individuals seeded in each cell/patch<br>(0: at demographic density dependence,<br>1: at half of the demographic density dependence,<br>2: according to quasi-equilibrium distribution) | 2 |

| Parameter | Description | Values |
|---|---|---|
| IndsHaCell | Specified number of individuals per hectare | 20 |
| PropStages | Proportion of initialised individuals in each stage | 0, 1, 0 |
| InitAge | Initial age distribution (0: minimum age for the respective stage, 1: random age between the minimum and maximum age for the respective stage, 2: according to a quasi-equilibrium distribution) | 2 |
| InitFreezeYear | Year until which species is confined to its initial range limits | 0 |
| RestrictRows | Number of rows at northern front to restrict range. range is restricted to northern front. | 0 |
| FinalFreezeYear | The year after which species is confined to its new, current range limits, after a period of range expansion. | 0 |

### Insect case study

#### Simulation parameters

| Parameter | Description | Values |
|---|---|---|
| Year | Number of simulated years | 50 |
| Replicates | Number of simulation iterations | 3 |
| Absorbing | Whether non-valid cells lead to direct mortality of the individual during transfer | FALSE |
| LocalExt | Local extinction | FALSE |
| EnvStoch | Environmental stochasticity (0: none, 1: global, 2: local) stochasticity acts (0: growth rate/fecundity, 1: demographic density dependence) | 0 |
| OutIntRange | Output of range file | 1 |

| Parameter | Description | Values |
|---|---|---|
| OutIntOcc | Output of occupancy file | 0 |
| OutIntPop | Output of population file | 0 |
| OutIntInd | Output of individual file | 1 |
| OutIntConn | Output of connectivity file | 0 |
| OutIntPaths | Output of SMS paths file | 0 |
| OutStartPop | Starting year for output population file | 0 |
| OutStartInd | Starting year for output individual file | 49 |
| OutStartConn | Starting year for output connectivity file | 0 |
| OutStartPaths | Starting year for output SMS paths file | 0 |
| SMSHeatMap | Output SMS heat map raster file | FALSE |

Landscape parameters

| Parameter | Description | Values |
|---|---|---|
| Landscape file | Filename(s) of the landscape map(s) | |
| Resolution | Resolution in meters | 80 |
| HabPercent | Whether habitat types/codes or habitat cover/quality | FALSE |
| NHabitats | Number of different habitat codes | 2 |
| K_or_DensDep | Demographic density dependence | 0, 44 |
| PatchFile | Filename(s) of the patch map(s) | |
| CostsFile | Filename(s) of the SMS cost map(s) | |
| DynamicLandYears | Years of landscape changes | 0 |
| SpDistFile | Filename of the species initial distribution map | |

Demography parameters

| Parameter | Description | Values |
|---|---|---|
| Rmax | Maximum growth rate<br>(number of offspring per female at very low density) | 1.39 |
| bc | Competition coefficient<br>(describes the type of density regulation) | 1 |
| ReproductionType | Decribes the reproduction type | 0 |

| Parameter | Description | Values |
|---|---|---|
| | (0: asexual/only female;<br>1: simple sexual model;<br>2: sexual model with explicit mating system) | |

Dispersal parameters

| Process | Parameter | Description | Values |
|---|---|---|---|
| Emigration | EmigProb | Emigration probabilities/parameters for each stage/sex | [0.4 1 5] |
| | SexDep | Sex-dependent emigration probability? | FALSE |
| | StageDep | Stage-dependent emigration probability? | FALSE |
| | DensDep | Density-dependent emigration probability? | TRUE |
| | UseFullKern | Shall the emigration probability be derived from dispersal kernel? | FALSE |
| Transfer | Type | Type of transfer | StochMove |
| | PR | Perceptual range | 1 |
| | PRMethod | Method to evaluate the effective cost<br>of a particular step from the landdscape<br>within the perceptual range:<br>1: Arithmetic mean<br>2: Harmonic mean<br>Weighted arithmetic mean | 2 |
| | MemSize | Size of memory, given as the number of previous steps<br>over which to calculate current direction<br>to apply directional persistence | 1 |
| | DP | Directional persistence. Corresponds to the<br>tendency to follow a correlated random walk. | 5 |
| | GoalType | Goal bias type<br>0: None, 2: dispersal bias | 0 |
| | GoalBias | Goal bias strength | 1 |

| Process | Parameter | Description | Values |
|---|---|---|---|
| | AlphaDB | Dispersal bias decay rate | -9 |
| | BetaDB | Dispersal bias decay inflection point (number of steps) | -9 |
| | Costs | Describes the landscapes resistance to movement: Either habitat-specific costs for each habitat type or ‘file’, to indictae to use the cost raster map(s) specified in the landscape module. | 100, 1 |
| | StepMort | Per-step mortality probability: constant or habitat-specific | 0.027 |
| | StraightenPath | Straighten path after decision not to settle in a patch? | TRUE |
| Settlement | DensDep | Density-dependent settlement requirements? | TRUE |
| | SexDep | Sex-dependent settlement requirements? | FALSE |
| | StageDep | Stage-dependent settlement requirements? | FALSE |
| | Settle | Settlement probability parameters for all stages/sexes. | $[1 \quad -10 \quad 1]$ |
| | MinSteps | Minimum number of steps | 0 |
| | MaxSteps | Maximum number of steps | 0 |
| | MaxStepsYear | Maximum number of steps per year | 0 |
| | FindMate | Mating requirements to settle? | FALSE |

#### Initialisation parameters

| Parameter | Description | Values |
|---|---|---|
| InitType | Type of initialisation<br>(0: free initialisation according to habitat map,<br>1: from loaded species distribution map,<br>2: from initial individuals list file) | 0 |
| FreeType | Option for free initialisation<br>(0: random in given number of cells,<br>1: all suitable cells/patches) | 1 |

| Parameter | Description | Values |
|---|---|---|
| | species distribution map (0: all suitable cells within all distribution presence cells, 1: all suitable cells within given number of randomly chosen presence cells) | |
| InitDens | Number of individuals seeded in each cell/patch (0: at demographic density dependence, 1: at half of the demographic density dependence, 2: according to quasi-equilibrium distribution) | 2 |
| IndsHaCell | Specified number of individuals per hectare | 20 |
| PropStages | Proportion of initialised individuals in each stage | 0 |
| InitAge | Initial age distribution (0: minimum age for the respective stage, 1: random age between the minimum and maximum age for the respective stage, 2: according to a quasi-equilibrium distribution) | 2 |
| InitFreezeYear | Year until which species is confined to its initial range limits | 0 |
| RestrictRows | Number of rows at northern front to restrict range. range is restricted to northern front. | 0 |
| FinalFreezeYear | The year after which species is confined to its new, current range limits, after a period of range expansion. | 0 |

#### Mammal case study

##### Simulation parameters

| Parameter | Description | Values |
|---|---|---|
| Year | Number of simulated years | 50 |
| Replicates | Number of simulation iterations | 3 |

| Parameter | Description | Values |
|---|---|---|
| Absorbing | Whether non-valid cells lead to direct mortality of the individual during transfer | FALSE |
| LocalExt | Local extinction | FALSE |
| EnvStoch | Environmental stochasticity (0: none, 1: global, 2: local) stochasticity acts (0: growth rate/fecundity, 1: demographic density dependence) | 0 |
| OutIntRange | Output of range file | 1 |
| OutIntOcc | Output of occupancy file | 0 |
| OutIntPop | Output of population file | 0 |
| OutIntInd | Output of individual file | 1 |
| OutIntConn | Output of connectivity file | 0 |
| OutIntPaths | Output of SMS paths file | 0 |
| OutStartPop | Starting year for output population file | 0 |
| OutStartInd | Starting year for output individual file | 49 |
| OutStartConn | Starting year for output connectivity file | 0 |
| OutStartPaths | Starting year for output SMS paths file | 0 |
| SMSHeatMap | Output SMS heat map raster file | FALSE |

Landscape parameters

| Parameter | Description | Values |
|---|---|---|
| Landscape file | Filename(s) of the landscape map(s) | |
| Resolution | Resolution in meters | 35000 |
| HabPercent | Whether habitat types/codes or habitat cover/quality | FALSE |
| NHabitats | Number of different habitat codes | 2 |
| K_or_DensDep | Demographic density dependence | 0, 0.000285 |
| PatchFile | Filename(s) of the patch map(s) | |
| CostsFile | Filename(s) of the SMS cost map(s) | |
| DynamicLandYears | Years of landscape changes | 0 |
| SpDistFile | Filename of the species initial distribution map | |

Demography parameters

| Parameter | Description | Values |
|---|---|---|
| Stages | Number of life stages | 4 |
| TransMatrix | Transition matrix. Defines the development probabilities from each stage into the next as well as the respective survival probabilities and fecundities | $\begin{bmatrix} 0 & 0 & 0 & 5 \\ 1 & 0 & 0 & 0 \\ 0 & 0.53 & 0 & 0 \\ 0 & 0 & 0.63 & 0.8 \end{bmatrix}$ |
| MaxAge | Maximum age in years | 17 |
| MinAge | Ages which an individual in stage i-1 must already have reached before it can develop into the next stage i. | 0, 0, 0, 0 |
| RepSeasons | Number of potential reproduction events per year | 1 |
| RepInterval | Number of reproductive seasons which must be missed following a reproduction attempt before another reproduction attempt may occur | 0 |
| PRep | Probability of reproducing in subsequent reproductive seasons | 1 |
| SurvSched | Scheduling of survival (0: at reproduction, 1: between reproductive events, 2: annually) | 1 |
| FecDensDep | whether density dependent fecundity probability is modelled | TRUE |
| DevDensDep | Whether density dependent development probability is modelled | FALSE |
| SurvDensDep | Whether density dependent survival probability is modelled | FALSE |

| Parameter | Description | Values |
| --- | --- | --- |
| DevDensCoeff | Relative density dependence coefficient for development | 1 |
| SurvDensCoeff | Relative density dependence coefficient for survival | 1 |
| FecStageWtsMatrix | Stage-dependent weights in density dependence of fecundity | Not selected. |
| DevStageWtsMatrix | Stage-dependent weights in density dependence of development | Not selected. |
| SurvStageWtsMatrix | Stage dependent weights in density dependence of survival | Not selected. |
| PostDestrictn | Whether individuals of a population die (FALSE) or disperse (TRUE) if its patch gets destroyed | FALSE |
| ReproductionType | Decribes the reproduction type (0: asexual/only female; 1: simple sexual model; 2: sexual model with explicit mating system) | 1 |
| PropMales | Proportion of males in the population | 0.5 |

Dispersal parameters

| Process | Parameter | Description | Values |
| --- | --- | --- | --- |
| Emigration | EmigProb | Emigration probabilities/parameters for each stage/sex | $\begin{bmatrix} 0 & 0 & 0.4 & 10 & 1 \\ 0 & 1 & 0.9 & 10 & 1 \\ 1 & 0 & 0 & 0 & 0 \\ 1 & 1 & 0 & 0 & 0 \\ 2 & 0 & 0 & 0 & 0 \\ 2 & 1 & 0 & 0 & 0 \\ 3 & 0 & 0 & 0 & 0 \\ 3 & 1 & 0 & 0 & 0 \end{bmatrix}$ |
| | SexDep | Sex-dependent emigration probability? | TRUE |
| | StageDep | Stage-dependent emigration probability? | TRUE |
| | DensDep | Density-dependent emigration probability? | TRUE |
| | UseFullKern | Shall the emigration probability be derived from dispersal kernel? | FALSE |

| Process | Parameter | Description | Values |
|---|---|---|---|
| Transfer | Type | Type of transfer | StochMove |
| | PR | Perceptual range | 1 |
| | PRMethod | Method to evaluate the effective cost<br>of a particular step from the landdscape<br>within the perceptual range:<br>1: Arithmetic mean<br>2: Harmonic mean<br>Weighted arithmetic mean | 2 |
| | MemSize | Size of memory, given as the number of previous steps<br>over which to calculate current direction<br>to apply directional persistence | 1 |
| | DP | Directional persistence. Corresponds to the<br>tendency to follow a correlated random walk. | 1 |
| | GoalType | Goal bias type<br>0: None, 2: dispersal bias | 0 |
| | GoalBias | Goal bias strength | 1 |
| | AlphaDB | Dispersal bias decay rate | -9 |
| | BetaDB | Dispersal bias decay inflection point (number of steps) | -9 |
| | Costs | Describes the landscapes resistance to movement:<br>Either habitat-specific costs for each habitat type<br>or ‘file’, to indictae to use the cost raster map(s)<br>specified in the landscape module. | 1000, 1 |
| | StepMort | Per-step mortality probability: constant or habitat-specific | 5e-04, 0 |

| Process | Parameter | Description | Values |
|---|---|---|---|
| | StraightenPath | Straighten path after decision not to settle in a patch? | TRUE |
| Settlement | DensDep | Density-dependent settlement requirements? | TRUE |
| | SexDep | Sex-dependent settlement requirements? | TRUE |
| | StageDep | Stage-dependent settlement requirements? | FALSE |
| | Settle | Settlement probability parameters for all stages/sexes. | $\begin{bmatrix} 0 & 1 & -10 & 1 \\ 1 & 1 & -10 & 1 \end{bmatrix}$ |
| | MinSteps | Minimum number of steps | 0 |
| | MaxSteps | Maximum number of steps | 14 |
| | MaxStepsYear | Maximum number of steps per year | 0 |
| | FindMate | Mating requirements to settle? | FALSE, TRUE |

Initialisation parameters

| Parameter | Description | Values |
|---|---|---|
| InitType | Type of initialisation<br>(0: free initialisation according to habitat map,<br>1: from loaded species distribution map,<br>2: from initial individuals list file) | 0 |
| FreeType | Option for free initialisation<br>(0: random in given number of cells,<br>1: all suitable cells/patches)<br>species distribution map (0: all suitable cells within<br>all distribution presence cells,<br>1: all suitable cells within given<br>number of randomly chosen presence cells) | 1 |
| InitDens | Number of individuals seeded in each cell/patch<br>(0: at demographic density dependence,<br>1: at half of the demographic density dependence,<br>2: according to quasi-equilibrium distribution) | 1 |

| Parameter | Description | Values |
|---|---|---|
| PropStages | Proportion of initialised individuals in each stage | 0, 1, 0, 0 |
| InitAge | Initial age distribution (0: minimum age for the respective stage, 1: random age between the minimum and maximum age for the respective stage, 2: according to a quasi-equilibrium distribution) | 2 |
| InitFreezeYear | Year until which species is confined to its initial range limits | 0 |
| RestrictRows | Number of rows at northern front to restrict range. range is restricted to northern front. | 0 |
| FinalFreezeYear | The year after which species is confined to its new, current range limits, after a period of range expansion. | 0 |

#### Reptile case study

##### Simulation parameters

| Parameter | Description | Values |
|---|---|---|
| Year | Number of simulated years | 50 |
| Replicates | Number of simulation iterations | 3 |
| Absorbing | Whether non-valid cells lead to direct mortality of the individual during transfer | FALSE |
| LocalExt | Local extinction | TRUE |
| LocalEctProb | Local extinction probability | 0.003 |
| EnvStoch | Environmental stochasticity (0: none, 1: global, 2: local) stochasticity acts (0: growth rate/fecundity, 1: demographic density dependence) | 0 |
| OutIntRange | Output of range file | 1 |
| OutIntOcc | Output of occupancy file | 0 |

| Parameter | Description | Values |
|---|---|---|
| OutIntPop | Output of population file | 0 |
| OutIntInd | Output of individual file | 1 |
| OutIntConn | Output of connectivity file | 0 |
| OutIntPaths | Output of SMS paths file | 0 |
| OutStartPop | Starting year for output population file | 0 |
| OutStartInd | Starting year for output individual file | 49 |
| OutStartConn | Starting year for output connectivity file | 0 |
| OutStartPaths | Starting year for output SMS paths file | 0 |
| SMSHeatMap | Output SMS heat map raster file | FALSE |

Landscape parameters

| Parameter | Description | Values |
|---|---|---|
| Landscape file | Filename(s) of the landscape map(s) | |
| Resolution | Resolution in meters | 15 |
| HabPercent | Whether habitat types/codes or habitat cover/quality | FALSE |
| NHabitats | Number of different habitat codes | 2 |
| K_or_DensDep | Demographic density dependence | 0, 1200 |
| PatchFile | Filename(s) of the patch map(s) | |
| CostsFile | Filename(s) of the SMS cost map(s) | |
| DynamicLandYears | Years of landscape changes | 0 |
| SpDistFile | Filename of the species initial distribution map | |

Demography parameters

| Parameter | Description | Values |
|---|---|---|
| Stages | Number of life stages | 3 |
| TransMatrix | Transition matrix.<br><br>Defines the development probabilities | $\begin{bmatrix} 0 & 0 & 0 & 0 & 1 & 12 \\ 1 & 0 & 0 & 0 & 0 & 0 \\ 0 & 1 & 0 & 0 & 0 & 0 \\ 0 & 0 & 0.21 & 0 & 0.68 & 0 \\ 0 & 0 & 0 & 0.28 & 0 & 0.41 \end{bmatrix}$ |

| Parameter | Description | Values |
|---|---|---|
| | from each stage into the next as well as the respective survival probabilities and fecundities | |
| MaxAge | Maximum age in years | 10 |
| MinAge | Ages which an individual in stage i-1 must already have reached before it can develop into the next stage i. | 0, 0, 0, 0, 0 |
| RepSeasons | Number of potential reproduction events per year | 1 |
| RepInterval | Number of reproductive seasons which must be missed following a reproduction attempt before another reproduction attempt may occur | 0 |
| PRep | Probability of reproducing in subsequent reproductive seasons | 1 |
| SurvSched | Scheduling of survival (0: at reproduction, 1: between reproductive events, 2: annually) | 1 |
| FecDensDep | whether density dependent fecundity probability is modelled | TRUE |
| DevDensDep | Whether density dependent development probability is modelled | FALSE |
| SurvDensDep | Whether density dependent survival probability is modelled | FALSE |
| DevDensCoeff | Relative density dependence coefficient for development | 1 |
| SurvDensCoeff | Relative density dependence | 1 |

| Parameter | Description | Values |
|---|---|---|
| | coefficient for survival | |
| FecStageWtsMatrix | Stage-dependent weights in density dependence of fecundity | Not selected. |
| DevStageWtsMatrix | Stage-dependent weights in density dependence of development | Not selected. |
| SurvStageWtsMatrix | Stage dependent weights in density dependence of survival | Not selected. |
| PostDestrictn | Whether individuals of a population die (FALSE) or disperse (TRUE) if its patch gets destroyed | FALSE |
| ReproductionType | Decribes the reproduction type (0: asexual/only female; 1: simple sexual model; 2: sexual model with explicit mating system) | 2 |
| PropMales | Proportion of males in the population | 0.5 |
| Harem | Maximum harem size | 3 |

Dispersal parameters

| Process | Parameter | Description | Values |
|---|---|---|---|
| Emigration | EmigProb | Emigration probabilities/parameters for each stage/sex | $\begin{bmatrix} 0 & 0 & 0.8 & 10 & 0 \\ 0 & 1 & 0.9 & 10 & 0 \\ 1 & 0 & 0.7 & 10 & 1 \\ 1 & 1 & 0.8 & 10 & 1 \\ 2 & 0 & 0.1 & 10 & 1 \\ 2 & 1 & 0.2 & 10 & 1 \end{bmatrix}$ |
| | SexDep | Sex-dependent emigration probability? | TRUE |
| | StageDep | Stage-dependent emigration probability? | TRUE |
| | DensDep | Density-dependent emigration probability? | TRUE |
| | UseFullKern | Shall the emigration probability be derived from dispersal kernel? | FALSE |
| Transfer | Type | Type of transfer | StochMove |

| Process | Parameter | Description | Values |
|---|---|---|---|
| | PR | Perceptual range | 1 |
| | PRMethod | Method to evaluate the effective cost of a particular step from the landdscape within the perceptual range:<br>1: Arithmetic mean<br>2: Harmonic mean<br>Weighted arithmetic mean | 2 |
| | MemSize | Size of memory, given as the number of previous steps<br>over which to calculate current direction<br>to apply directional persistence | 1 |
| | DP | Directional persistence. Corresponds to the tendency to follow a correlated random walk. | 5 |
| | GoalType | Goal bias type<br>0: None, 2: dispersal bias | 0 |
| | GoalBias | Goal bias strength | 1 |
| | AlphaDB | Dispersal bias decay rate | -9 |
| | BetaDB | Dispersal bias decay inflection point (number of steps) | -9 |
| | Costs | Describes the landscapes resistance to movement:<br>Either habitat-specific costs for each habitat type<br>or ‘file’, to indictae to use the cost raster map(s)<br>specified in the landscape module. | 10, 1 |
| | StepMort | Per-step mortality probability: constant or habitat-specific | 0.05, 0 |
| | StraightenPath | Straighten path after decision not to settle in a patch? | TRUE |

| Process | Parameter | Description | Values |
| --- | --- | --- | --- |
| Settlement | DensDep | Density-dependent settlement requirements? | TRUE |
| | SexDep | Sex-dependent settlement requirements? | FALSE |
| | StageDep | Stage-dependent settlement requirements? | FALSE |
| | Settle | Settlement probability parameters for all stages/sexes. | $[1 \quad -10 \quad 0.7]$ |
| | MinSteps | Minimum number of steps | 0 |
| | MaxSteps | Maximum number of steps | 420 |
| | MaxStepsYear | Maximum number of steps per year | 40 |
| | FindMate | Mating requirements to settle? | FALSE |

Initialisation parameters

| Parameter | Description | Values |
| --- | --- | --- |
| InitType | Type of initialisation<br>(0: free initialisation according to habitat map,<br>1: from loaded species distribution map,<br>2: from initial individuals list file) | 0 |
| FreeType | Option for free initialisation<br>(0: random in given number of cells,<br>1: all suitable cells/patches)<br>species distribution map (0: all suitable cells within all distribution presence cells,<br>1: all suitable cells within given number of randomly chosen presence cells) | 1 |
| InitDens | Number of individuals seeded in each cell/patch<br>(0: at demographic density dependence,<br>1: at half of the demographic density dependence,<br>2: according to quasi-equilibrium distribution) | 2 |
| IndsHaCell | Specified number of individuals per hectare | 20 |
| PropStages | Proportion of initialised individuals in each stage | 0, 1, 0 |
| InitAge | Initial age distribution | 2 |

| Parameter | Description | Values |
| --- | --- | --- |
| | (0: minimum age for the respective stage, 1: random age between the minimum and maximum age for the respective stage, 2: according to a quasi-equilibrium distribution) | |
| InitFreezeYear | Year until which species is confined to its initial range limits | 0 |
| RestrictRows | Number of rows at northern front to restrict range. range is restricted to northern front. | 0 |
| FinalFreezeYear | The year after which species is confined to its new, current range limits, after a period of range expansion. | 0 |
